# Unveiling the Terra Cognita of Sequence Spaces using Cartesian Projection of Asymmetric Distances

**DOI:** 10.1101/2025.09.04.674223

**Authors:** Alban Ramette

**Affiliations:** Institute for Infectious Diseases, University of Bern, Bern, Switzerland; Multidisciplinary Center for Infectious Diseases, University of Bern, Bern, Switzerland

**Keywords:** sequence analysis, multiple sequence alignment, genetic distances

## Abstract

Visualizing relationships within massive biological datasets remains a significant challenge, particularly as sequence length and volume increase. We introduce CAPASYDIS (Cartesian Projections of Asymmetric Distances), a scalable approach designed to map the explored regions of a given sequence space. Unlike traditional dimensionality reduction methods, CAPASYDIS calculates asymmetric distances which account for both the position and type of sequence variations. It projects sequences into a fixed, low-dimensional coordinate system, termed a “seqverse”, where each sequence occupies a permanent location. This design allows for the instant mapping of new sequences without the need to recalculate the global structure, transforming sequence analysis from a relative comparison into navigation on a standardized map. We applied this method to a large rRNA sequence dataset spanning the three domains of life. Our results demonstrate that the sequences of Bacteria, Archaea, and Eukaryota occupy spatially distinct regions characterized by fundamentally different shapes and patterns of variation. Furthermore, the resulting seqverses retain high amount of taxonomic information, when analyzed from broad domain levels to single-base differences. Overall, CAPASYDIS provides a reproducible, scalable framework for defining the boundaries and topography of biological sequence universes.

## Introduction

Genetic diversity analysis and phylogenetic reconstruction are cornerstones of biological research. They provide insights into evolutionary relationships, ecological dynamics, and the intricate workings of life itself. The advent of next-generation sequencing (NGS) has unveiled the vast extent and complexity of genetic diversity, revealing that even few grams of soil (1), milliliters of sea water (2), or square centimeters of human skin (3), can harbor a staggering array of lifeforms. The unprecedented availability of sequence data presents opportunities, but also challenges for data analysis and visualization (4). Traditional methods, such as distance matrix calculations and phylogenetic tree reconstruction, provide essential insights into genetic diversity. However, these methods often struggle to efficiently process and visualize the high-dimensional complexity of sequence space (5, 6), particularly as datasets grow in scale. A primary limitation is that these techniques do not provide a fixed coordinate system for sequences in a low-dimensional space; instead, they rely on heuristics to approximate the positioning of sequence points within projections or dendrograms, as explained below.

If we were to represent a complete map of all possible sequence variations for a given sequence, the curse of dimensionality would make it computationally infeasible to compare all possible sequence combinations. For a DNA sequence of length n, there are 4^n^ possible sequence variations. The corresponding distance matrix between all those sequences would then be a matrix of size 4^2n^. As sequence length grows, this number increases exponentially, quickly exceeding the computational limits of even the most powerful supercomputers. Even for even a 1,000-base sequence, an exhaustive map of all possible evolutionary possibilities is impossible to create, as it corresponds approximately to 10^1,204^ possible sequence combinations. In comparison, the estimated total number of stars in an inflationary universe (observed and unobserved) would be in the range of 10^100^ (7). This challenge forces researchers to rely on methods that only compare the sequences actually observed in their datasets, rather than representing the entire potential sequence space. Computational approaches, such as dimensionality reduction, may be used to represent large distance matrices in a reduced space (8). Yet, they inevitably sacrifice some information (variance, topology, or global structure) to reduce the complexity of the data, as observed in Principal Component Analysis (PCA)(9), t-Distributed Stochastic Neighbor Embedding (t-SNE) (10, 11), or Uniform Manifold Approximation and Projection (UMAP)(12), to cite a few. Furthermore, applying these techniques to massive sequence datasets presents significant scalability challenges, as methods that rely on all-vs-all distance matrices become computationally prohibitive as datasets grow. Additionally, many non-linear methods (like t-SNE) lack a native parametric mapping, meaning that projecting new sequences often requires re-calculating the entire embedding rather than navigating a fixed, existing map.

Beyond the computational burden of generating comprehensive distance matrices, the construction of evolutionary trees, such as dendrograms or phylograms, presents itself a significant independent challenge. Even for a modest number of observed sequences, the number of possible tree topologies grows at a rate that is superexponential. It means that this number increases far faster than an exponential function: For example, the number of possible unrooted binary trees for just 10 sequences is over two millions, and it is a number with 74 digits for 50 sequences (5, 13). This consideration makes it impractical to search for a single, “perfect” tree. As a result, bioinformatic methods for tree construction must rely on heuristics, which are algorithms that use assumptions to find a reasonably good, though not necessarily optimal, solution. Approaches, such as Neighbor-Joining or Maximum Likelihood, iteratively build a tree that best fits the input data, but they do not guarantee that the absolute best tree has been found. This reliance on heuristics introduces a level of uncertainty and subjectivity into the analysis. Another key limitation lies in how dendrograms and phylograms are visualized. These diagrams represent tree topology (the branching pattern) and branch lengths (often proportional to evolutionary distance). Even though they are powerful methods for displaying hierarchical relationships among sequences, they are not meant to convey information about the specific spatial coordinates of sequences in the representation. This means that, while a tree can be visually modified by simply rotating a branch at any node, the underlying evolutionary relationships remain exactly the same. This non-uniqueness can create a misleading visual interpretation, especially when tree size grows larger, making it difficult to compare individual sequence positions across experiments or studies.

To address limitations in both scalability and visualization, we introduce CAPASYDIS (Cartesian Projections of Asymmetric Distances), a framework designed to map and visualize the “terra cognita” of a given sequence space. CAPASYDIS embraces the inherent asymmetry in reading direction of a pair of aligned sequences, and acknowledges that, even if the same changes between two sequences exist, if those changes occur at different sequence positions, they should have *different impacts on the resulting distances* between those two sequences. Starting from a multiple sequence alignment, the novel concept associates a distinct distance value to each unique query sequence relative to a given reference sequence. Then using those unique vectors, a reduced dimensional space can be built by plotting the sequences using those unique coordinates into a lower-dimensional space. By employing Cartesian projections, sequences are mapped onto a visualizable space with predefined, fixed axes, enabling reproductible exploration and analyses, because the coordinates of the mapped sequences do not change as the dataset grows larger. In this study, the CAPASYDIS methodology is presented and illustrated by analyzing a large dataset (rRNA gene sequences) where the known diversity of the three domains of life is investigated at different taxonomic levels.

## Methods

### Data set

The file “SILVA_138.2_SSURef_NR99_tax_silva_full_align_trunc.fasta.gz” was downloaded from the SILVA rRNA database project (https://www.arb-silva.de; (14, 15)) on 2024-07-11. The dataset is based on the full SSU Ref 138.2 dataset and include a total of 510,508 rRNA sequences from the three domains of life. The CAPASYDIS processed data used for the NR99 analysis is available at https://doi.org/10.5281/zenodo.17055026.

### Implementation

The initial proof-of-concept was implemented with the R language (version 4.4.1; https://www.r-project.org/) to develop the algorithm using small datasets. To allow the pre-processing of large Multiple Sequence Alignments (MSA), further computational scripts were developed in the *go* language (aka “Golang”; https://go.dev/; version 1.24.5) based on the *biogo* library (version v1.0.1; (16)). Further analyses of the csv files produced by CAPASYDIS *go* scripts were processed with the R library *dplyr* (version 1.1.4; (17)), and visualized using the R libraries *ggplot2* (version 3.5.2; (18)), *plotly* (version 4.10.4; (19)), *grid* (version 4.4.1; base R), *geometry* (version 0.5.2; (17)), *FNN* (version 1.1.4.1; https://cran.r-project.org/web/packages/FNN/), *patchwork* (version 1.3.0; https://cran.r-project.org/web/packages/patchwork/index.html), *ggalign* (version 1.0.2; (20)), and *forcats* (version 1.0.0; https://cran.r-project.org/web/packages/forcats/index.html). Interactive data visualizations were also developed using the JavaScript libraries *D3.js* (version 7.9.0; https://d3js.org/) and *Three.js* (version 0.179.1; https://threejs.org/), enabling the direct rendering of large-scale objects within the user’s web browser. Gemini 2.5 Flash and Pro were used throughout the project to help develop and correct code snippets in the R, Golang and JavaScript languages. All CAPASYDIS analyses on NR99 datasets were performed using an AlmaLinux 8.10 compute server (AMD EPYC-Rome processors 2.26 GHz; 491 GB of RAM). Further data analyses in R or with the JavaScript libraries were done on a standard Windows office laptop (Intel Core Ultra 5 processor 135U with 1.60 GHz, 16 GB RAM, 8 GB Intel Graphics GPU).

### Code availability

R, Golang and JavaScript scripts are available at https://github.com/RametteLab/CAPASYDIS. The repository also includes the workflow and scripts used to analyze the NR99 dataset presented in the study.

## Results

### Concepts

CAPASYDIS associates each query sequence in a MSA to a unique and fixed Euclidean coordinate. Given a sequence alignment of length *n* bases between a candidate sequence S and a reference sequence R, a *score xi* for the base at position *i* of sequence S is first calculated as:

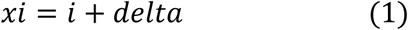

where *delta* is null, except if the base at position *i* is different between S and R. In the latter case, the delta value is either a small constant value (e.g. 0.01) as compared to the magnitude of the indices, as in the simple example (**Fig. 1A**), or a unique value associated with a specific base pair combination (see next section below; **Table 1**). Second, the individual scores are square-root transformed and then summed up across all the sequence positions:

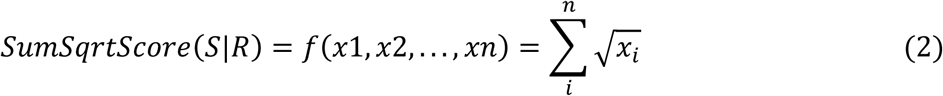

where *xi* represents the score at position *i* of the sequence S, defined in Equation 1.

**Figure 1.**
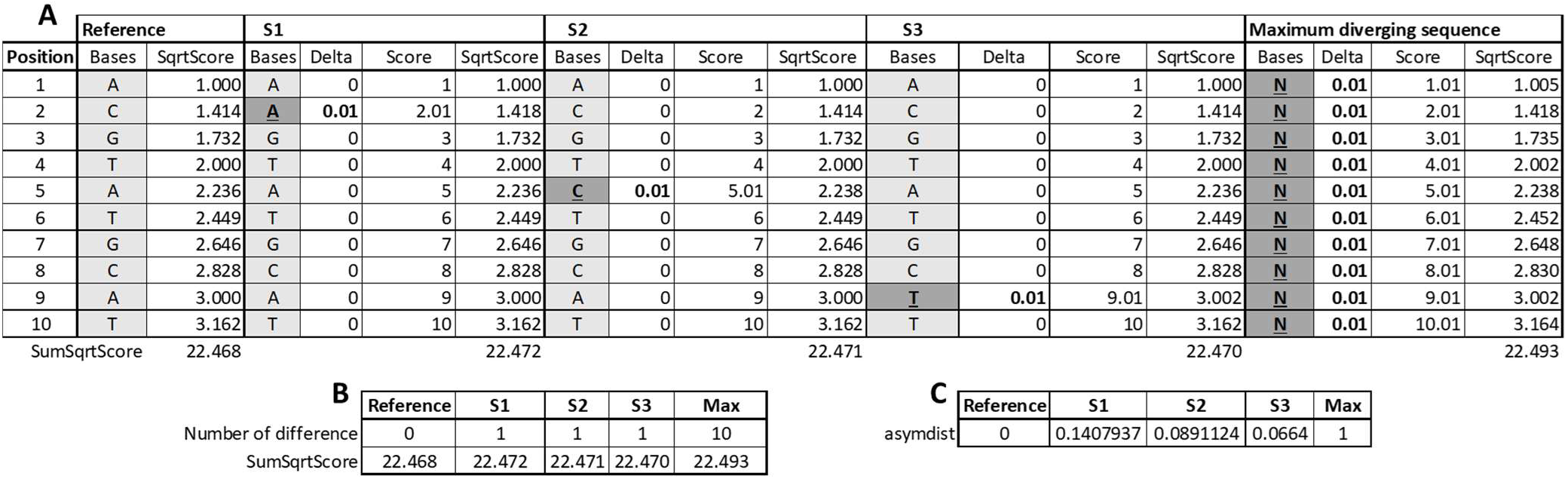
A conceptual example. **A**) Example of alignment dataset detailing the calculation of **B**) *SumSqrtScore* (Eq. E2), and **C**) *asymdist* (Eq. E3) consisting of re-scaled *SumSqrtScore* values to [0,1] range for the simple case of constant *delta* (0.01 for a base difference; 0 otherwise). Sequences S1, S2, and S3, differ only by a single mutation as compared to the reference sequence, at the beginning, middle and end of their sequences, respectively. The maximum diverging sequence contains “N“ (i.e. unknown, yet different base) at all positions of the aligned sequences.

**Table 1.** A possible coding of case-specific delta values.

| S\|R change | *delta* value |
| --- | --- |
| Transition |  |
| A ↔ G | 0.01 |
| C ↔ T | 0.02 |
| Transversion |  |
| A ↔ C | 0.06 |
| G ↔ T | 0.07 |
| A ↔ T | 0.08 |
| G ↔ C | 0.09 |
| Unknown |  |
| - ↔ A | 0.11 |
| - ↔ T | 0.12 |
| - ↔ G | 0.13 |
| - ↔ C | 0.14 |

For example, a one-base difference at position 2 between sequences S1 and R (both 10-base long) leads to the following computations for S1: SumSqrtScore (S1|R)= √1 + √2.01 + √3 + … + √10 = 22.472. For R, we obtain: SumSqrtScore (R|R)= √1 + √2 + √3 + … + √10 = 22.468 (**Fig. 1B**).

The function *SumSqrtScore* (Equation 2) produces a numeric sequence that diverges indefinitely as *n* increases through all positive integers. There is no simple, concise formula to directly calculate the n^th^ term of the sequence without performing the summation of square roots (no closed-form expression). Importantly, the gaps between successive *n* terms increase, which means that terms appearing later will create a larger effect on the overall distance for a given sequence. This is because the difference between consecutive terms [f(n + 1) – f(n)] is equal to √*n* + 1, and as *n* increases, so does the square root, meaning the gaps between terms get wider and wider. Third, the output of Equation 2 is then scaled to the interval [0,1]:

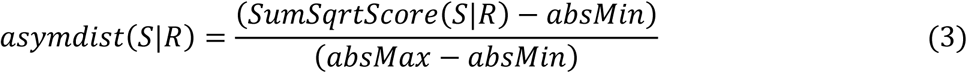

where *SumSqrtScore*(*S*|*R*) is calculated using Equation 2, *absMin* is the smallest value obtained if the S and R are identical sequences, and *absMax* is the largest value that is obtained if the sequence S differs from R at all positions using the maximal possible *delta* value*s*. Equation 3 provides a quantity, *asymdist*, which reflects the asymmetric nature of the results, as the calculation takes into account the positions at which base modifications occur along the alignment. The approach provides distinct results than the number of base differences, which stay constant in case of one mutation occurring at different position along the sequence (**Fig. 1B**). Changes that occur early in the sequences i.e. towards the 5’ end of the reference sequence, as illustrated by S1, have more impact (i.e. they modify the coordinate to a larger degree) than changes occurring later (i.e. towards the 3’ end, such as S3), and overall produce larger *asymdist* values (**Fig. 1C**). This reflects how CAPASYDIS is designed: The index value captures the positional information difference between the sequences and the reference, while the *delta* value adds a little, case-specific information, and both sources of information are combined into a mathematically unique value, owing to the property of the square-root function (Equation 2). For instance, the effect of *delta* on a small value, i.e. a small index (e.g. 1), √1 + 0.01 = 1.0049 and is to be compared to 1, when no difference between the two sequences at position 1 is observed. Using the same *delta* but at position 100 leads to an effect that is ten times smaller than that at position 1, given that √100 + 0.01 =10.00049, which is to be compared to 10 when no difference is observed at position 100.

The asymmetric nature of the proposed metric can be illustrated graphically using a set of 100 aligned sequences, which differ each by one base only to the reference sequence. By design, a distinct *asymdist* value is associated with each unique sequence provided that the sequence differs from the reference in a unique manner, as illustrated by the marginal histograms. After sorting sequences on the x axis by the position of their modification, from 5’ to 3’ (**Fig. 2A,B)**, a larger impact on the *asymdist* values is evidenced by changes that occur at the beginning than towards the ends of the sequences. In addition, classical phylogenetic distances could not discriminate among those 100 sequences that differ from the reference sequence by only one single variation (**Fig. 2C)**: The JC69 model produces only one constant value for all sequences, which is not surprising given that this model assumes that equal probability for a base change, regardless of its position along the sequence (21). Similar results are obtained if one relaxes the assumption of equal base frequencies, by using e.g. the F81 model, which is a generalized Jukes–Cantor model (22). Alternatively, one may also try probabilistic models for nucleotide substitution such as the BH87 model (23), which accounts for the observed proportions of changes among the four bases. This latter distance is not symmetric (**Fig. 2D**), but results in only two distance classes (0.00125, 0.00083) for all 100 distinct sequences. A similar result is obtained when using the TN93 model (24), which assumes distinct rates for both kinds of transitions (A ↔ G versus C ↔ T), and transversions: Here, the base frequencies are not assumed to be equal and are estimated from the data. The TN93 model produced three classes of values corresponding to distances of 0.01008, 0.01021, and 0.01021 (**Fig. 2E**). Overall, the *asymdist* calculation approach allows the set of 100 query sequences to be associated each with definitive and unique coordinates, as it integrates both the presence and the position of a change.

**Figure 2.**
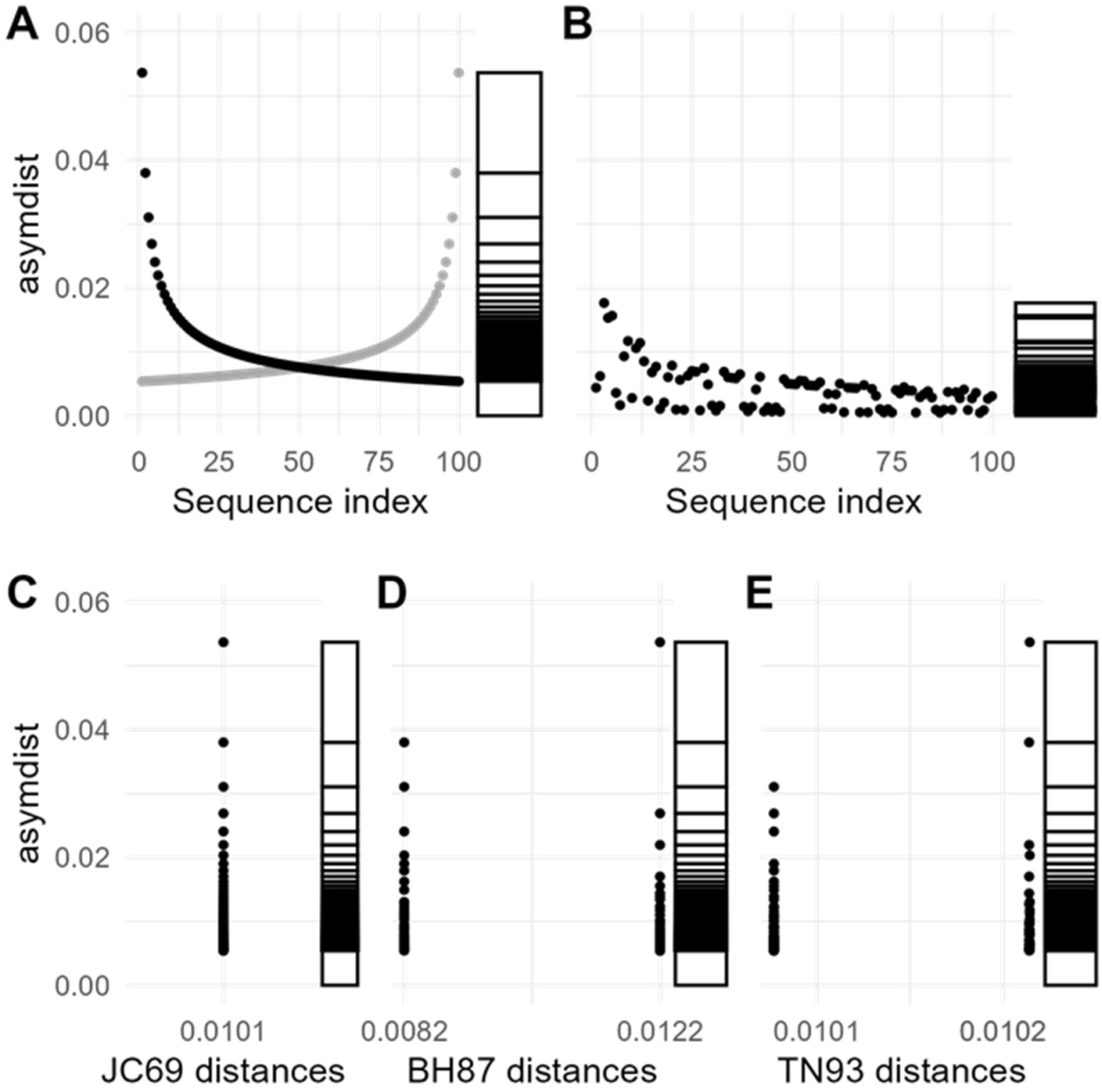
Illustration of *asymdist* properties on a set of 100 aligned sequences. A 100-base long nucleotide sequence was modified one base at the time, from left to right, to generate a set of 100 distinct sequences. **A**) These sequences were sorted by the position (index) of the nucleotide change in the sequences (x axis) from left to right, and *asymdist* with constant delta (0.01) was calculated for each sequence using the original sequence as reference. The y axis depicts the asymdist for each of those sequences (black dots), with a marginal histogram to confirm the uniqueness of each *asymdist* value. Grey dots correspond to asymdist calculated for the reverse order of the aligned sequences (reading from right to left). **B**) *asymdist* values were calculated using case-specific *delta* values, as proposed in **Table 1** for the same set of sequences. Comparison of *asymdist* values (*delta* = 0.01) with distances estimated by different genetic distance models: **C**) Jukes–Cantor 1969 (JC69) or Felsenstein 1981 (F81), **D**) Barry and Hartigan 1987 (BH87), and **E**) Tamura and Nei 1993 (TN93) models for the same set of 100 sequences.

### Using distinct *delta* values

When using a constant *delta* value in Equation 1, different types of changes at a given position would not be distinguishable from one another, and would produce asymdist coordinates that are thus mathematically identical. To create a unique numerical weight assigned to each specific change, distinct delta values for each possible mismatch, including those involving gaps in the reference sequence, can be proposed (**Table 1**). Those case-specific delta values further numerically modify the final asymdist score of the sequence based on both the position and nature of the change that occurred. When using case-specific delta values, the uniqueness of the coordinates of each single sequence is maintained (marginal histogram; **Fig. 2B**), but also that a different, more granular pattern is obtained for the set of 100 sequences as compared to using a unique delta value for all changes (**Fig. 2A**).

Although this coding of case-specific delta values is arbitrary, from a mathematical point of view, we can justify this choice by stating that delta values need to be relatively smaller than the smallest index values (delta << 1; 1 being the smallest index value in Equation 2). From a biological point of view, we chose to reflect common biological knowledge by assuming that transitions (A ↔ G, C ↔ T) involve bases of similar shape, and are thus less likely to result in amino acid substitutions, and are therefore more likely to persist as “silent substitutions”, and could thus be more likely to occur. On the contrary, transversions (A ↔ C, G ↔ T, A ↔ T, G ↔ C) involve exchange of one-ring and two-ring structures, and as such, transversions are generated at lower frequency than transition mutations. Probably other options of coding *delta* values are also suitable, as long as the magnitude of those values is kept smaller than the smallest index value.

### A possible workflow using CAPASYDIS

In essence, CAPASYDIS produces unique and definite coordinates for each sequence in a MSA. It is thus essential that the starting dataset consists of a set of high-quality sequences, for which the aligned positions are meaningful. As such, a series of preparatory steps may be needed before applying the CAPASYDIS approach to a MSA in order to build a 2D or 3D visualization of the sequence space, which we defined as “seqverse” (**Fig. 3**). We start with a MSA of rows of sequences by columns of aligned bases in which A, U, T, G, C, – (dash, or alignment gap) and “.” (missing base) are considered valid information. In step **a**) the MSA is vertically truncated to keep only the positions of the MSA that are not missing or not a constant gap for a given k proportion of bases (step a_1_). For instance, for k = 0.9, the MSA is truncated at specific starting and ending positions so that 90% of the sequences starting in the new MSA contain at least k bases that are not missing bases (“.”). The same approach is applied to the last position of the alignment. Positions consisting of “-” or “.” only, as well as duplicated sequences, are removed from the MSA in a second stage (a_2_). The headers of the remaining FASTA sequences are appended with the count of identical sequences found in the MSA. In step **b**) CAPASYDIS coordinates are computed for all sequences in the MSA based on the indices of the chosen reference sequences, and the 2D (or 3D) coordinates are merged into a single csv file. In step **c**) specific color labels can optionally be added to sequence features (e.g. color “red” for a given species or taxonomic level such as the domain “Archaea”). In step **d**) the coordinate file of the seqverse can be displayed in one to three dimensions using dynamic visualizations, which allow the interactive analysis of very large number of points and their patterns. The user can thus identify points interactively, delineate grouping structures (hulls, contours), and hover to see details, zoom to explore fine-grained topics, or retrieve the information from specific sequences. Optionally, identified sequences or patterns can be retrieved from the original MSA (step **e**). Those steps are illustrated with an example in the next section.

**Figure 3.**
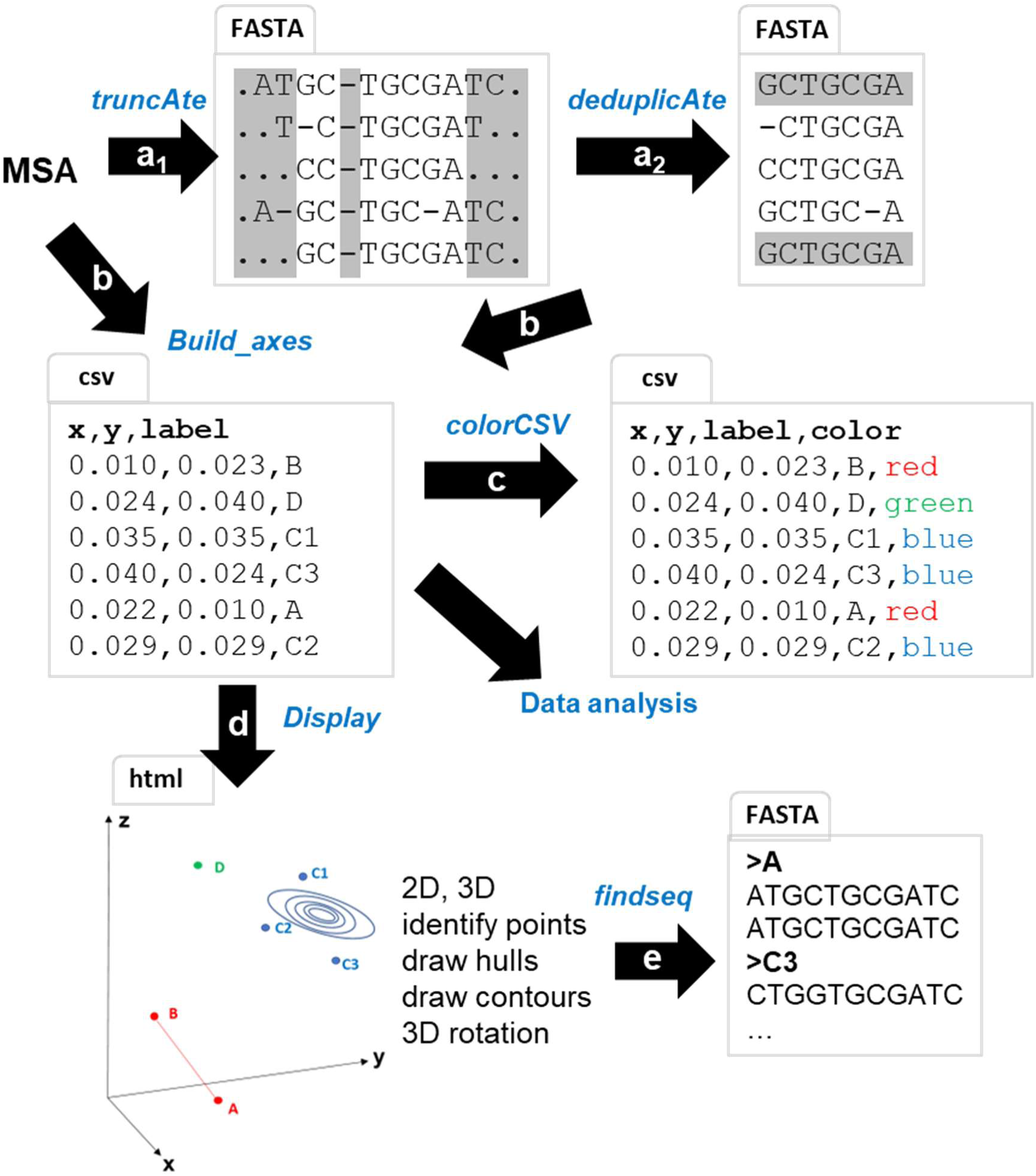
Workflow for CAPASYDIS. Starting from a multi-sequence alignment (MSA), the following steps are performed: **a**_1_) The MSA is truncated to keep only the positions of the MSA that are not missing or unknown bases for a given k proportion of bases. In grey, the information that is removed from the MSA. **a**_2_) Redundant sequences in the MSA are removed. **b**) the asymdist distances are computed relative to the chosen reference and are merged into one CSV file. Bad entries in the taxonomy fields are removed. **c**) Specific colors can be assigned to sequence features (e.g. taxonomic levels) to facilitate their visualization. The file can also be used directly for further data analyses. **d**) The coordinate file can be displayed interactively to allow exploration of the seqverse. **e**) Sequences can be retrieved from the original MSA for downstream processing. The resulting file formats (FASTA, csv or html) are also indicated for each step.

In practice, a crucial part of this workflow is the selection of the reference sequences that define the CAPASYDIS axes, and we outline here a few concrete guidelines for users. For the first reference, we recommend choosing a representative, abundant, and well-annotated sequence from the focal group (for example, a model organism or a taxonomically central lineage), so that the origin of the seqverse is anchored in a densely sampled and biologically well-understood region of sequence space. In our NR99 case study, we therefore selected the 16S rRNA gene of Escherichia coli O157:H7 as the first reference because it is both extensively characterized and among the most abundant *Escherichia–Shigella* sequences in the cleaned MSA. From this starting point, additional axes can be defined iteratively by using the *build_axes*() function to identify the most distant sequence (maximal asymdist) to the current reference and then using that sequence as the next reference. Depending on the analytical goal, users may either (i) construct global maps by starting from a widely used model taxon and iteratively adding the most distant sequences to capture maximal diversity, or (ii) construct local, taxon-focused maps by choosing all references from within the same clade to emphasize internal structure. Finally, we recommend verifying that the first reference has many closely related sequences, avoiding references with extensive missing or ambiguous data, and, if needed, testing alternative first references on a subset of the data to ensure that major qualitative patterns are robust.

### The rRNA seqverses of the three domains of life

The CAPASYDIS workflow was applied to ribosomal RNA sequences obtained from the SILVA release 138 SSU Ref NR99, which includes 510,495 RNA sequences with 20,389 (3.99%) sequences classified as Archaea, 58,940 (11.54%) as Eukaryota, and 431,166 (84.46%) as Bacteria. This large MSA consists of 50,000 aligned nucleotide positions, which was originally designed to accommodate sequence length variations and hypervariable regions within the 16S and 18S rRNA gene sequences, as an universal alignment profile (14). This stable framework is thus robust to the inclusion of new sequences (total information content of 25,524,750,000 bases), but may also contain many columns with no or little information. Thus, the script truncAte was first applied with a threshold k of 0.90 that select positions in the truncated MSA to be kept if 90% of the bases at those positions are not missing information or alignment gaps. The procedure also removes sequences with unknown (“N”) bases or IUPAC nucleotide codes. The analysis was completed in 1h16 min when using 60 cores and produced a new MSA of dimensions 387,633 sequences by 29,932 aligned base positions, so keeping effectively around 76% of the initial sequences, and 60% of the initial aligned base positions. The resulting MSA was further deduplicated and degapped with deduplicAteSeq, resulting in 331,663 unique sequences. This step took about 10 min using 30 cores. This deduplicated MSA now encompassed 6,031 sequences for Archaea (1.8%), 291,195 sequences for Bacteria (87.8%) and 34,437 sequences for Eukaryota (10.4%), resulting in a total dataset size of 9,927,336,916 bases, with average sequence lengths of 1,278 bases (minimum 842, maximum 2,787) for Archaea, 1,313 (1,036, 2,642) bases for Bacteria, and 1,605 (1,087, 3,232) bases for Eukaryota.

Escherichia coli O157:H7 (GenBank Accession number: AB035920; identified as AB035920.964.2505 in the MSA) was chosen as first reference for the CAPASYDIS axes, as its sequence was the most abundant among the Escherichia-Shigella sequences with a total of 379 sequences in the cleaned MSA. The function build_axes() provides several parameters such as “-stat” to retrieve sequence identity of the most distant sequences, the overall mean, standard deviation and maximum asymdist values, based on the input of the index of the chosen reference sequence. When applied to our example using case-specific delta values (**Table 1**), the most distant sequence to the 16S rRNA gene of Escherichia coli O157:H7 was the 18S rRNA gene of the marine eukaryotic cellular organism Heterostegina depressa (GenBank accession number AJ879131.1; NR99 accession AJ879131.1.3425). The latter is classified in the Phylum Foraminifera, and produced the largest asymdist value of 0.0983304. The second CAPASYDIS axis for the NR99 MSA was then calculated using as reference the H. depressa sequence in the MSA. The build_axes() function identified the 16S rRNA gene sequence of the archaeon Pyrobaculum sp. (GenBank accession number AB302407.1; NR99 accession AB302407.1.2962) at an asymdist value of 0.1298366, as the most distantly related to the Eukaryotic reference. Finally, the asymdist values with references to E. coli (Bacteria; axis “x”), H. depressa (Eukaryota; axis “y”), and Pyrobaculum (Archaea; axis “z”) were merged into a comma-separated-value (csv) file, with a simple structure of the type “x, y, z, label”, where (x, y, z) represents the asymdist coordinates in different dimensions, and the “label” describes the taxonomic information string of the sequences. asymdist calculation via build_axes() for one reference sequence in the MSA took about 1-2 min using 60 cores in parallel. The uniqueness of the coordinates of each sequence was obtained with a decimal point precision of e-10. The choice of the precision level (tunable parameter in the build_axes script) depends on the intrinsic properties of the MSA, such as dimensions and complexity. Smaller and simpler MSA would require less precision to obtain unique point coordinates for each sequence in the seqverse. For our dataset, the needed numerical precision was empirically determined by defining the number of decimals that can differentiate a sequence with only one minimal change (smallest delta value) at the most proximal index position.

The taxonomic information provided with the NR99 was further parsed to remove inconsistencies and missing information using the script SILVA_go, which is a Golang script that removes erroneous terms (e.g. habitat or sample descriptors) in the taxonomic fields. After removing taxonomic inconsistencies, the data set labels contained 14 phyla and 28 classes for Archaea, 92 phyla and 185 classes for Bacteria, while Eukaryota consisted of 10 phyla and 21 classes. Taxonomic levels in the coordinate file were further associated with specific color labels using either the R programming language or Golang (colorCSV). The resulting labelled coordinate file was finally rendered with the R library plotly (**Fig. 4**) or via a suite of HTML interactive pages using the JavaScript 3D (Three.js) or 2D (D3.js) libraries, which are suitable for rendering very high-dimensional data. Several exemplary JavaScript scripts are provided in the package repository.

**Figure 4.**
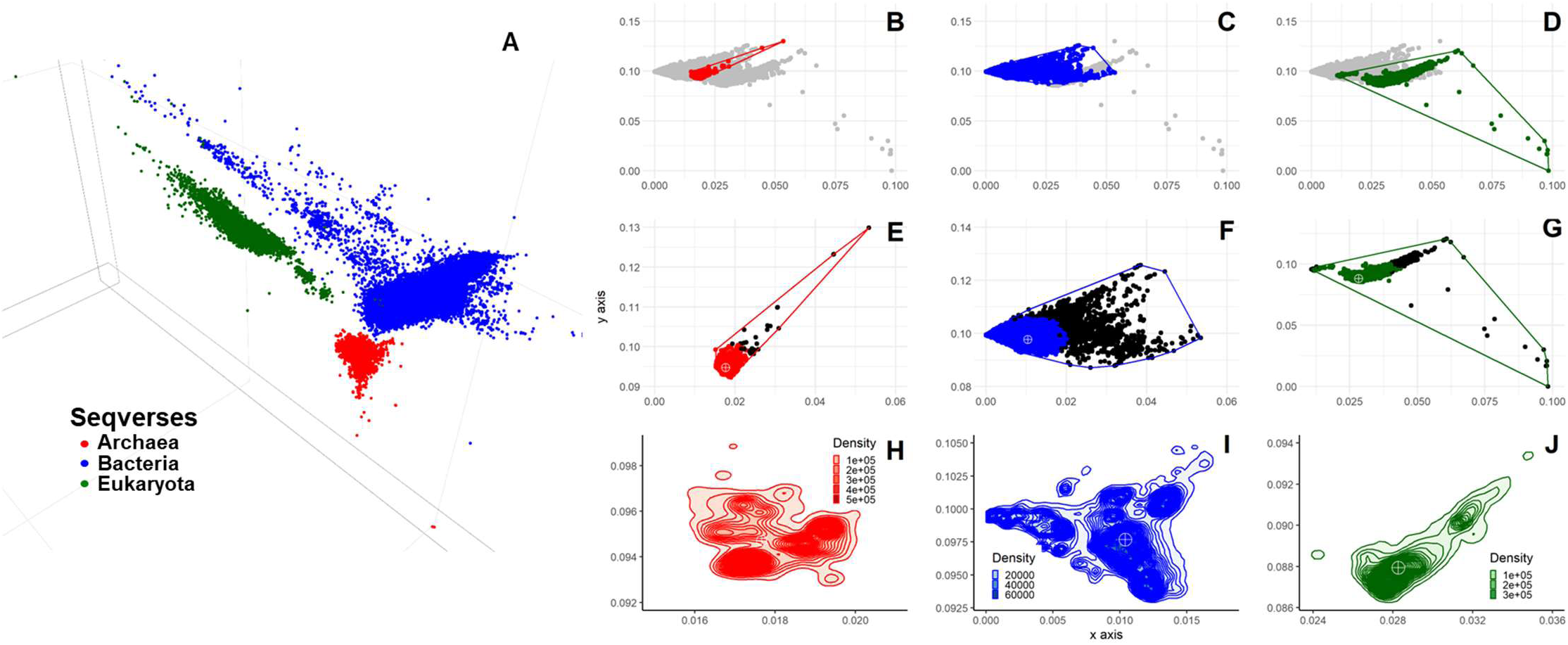
Visualization of the rRNA gene seqverse (n = 331,663 distinct sequences) of the three domains of life. The NR99 MSA was analyzed with the CAPASYDIS workflow, which produced *asymdist* values with references to *E. coli* (Bacteria; axis “x“), *H. depressa* (Eukaryota; axis “y”), and *Pyrobaculum* (Archaea; axis “z“). Each sequence point was colored by its associated domain with Archaea in red Bacteria in blue, Eukaryota in green. **A**) Three-dimensional visualization of the 16S rRNA gene seqverse). Two dimensional convex hulls of the **B**) Archaea seqverse, **C**) Bacteria seqverse, and **D**) Eukaryota seqverse, with sequences from other domains colored in grey. Panels **E)**, **F)**, **G)** represent magnified views of each respective seqverse with sequences belonging to the 1% most distant sequences to the domain centroid indicated in black points. Panels **H), I), J)** represent 2D kernel density estimations (with 50 bins) which are displayed as contour lines, for Archaea, Bacteria, and Eukaryota, respectively. The domain centroids are indicated as white markers in panels **E** to **J**.

The rRNA gene seqverses of the three domains of life fall into distinct, non-overlapping regions (**Fig. 4A**), and spanned different sizes and shapes (**Fig. 4B-J**). The 3D projection of the seqverses indicated that the Eukaryota seqverse appears rather flat, that of the Archaea compact and conic, while the bacterial seqverse seems to display an extensive “wing” shape (**Movies 1, 2**). Two-dimensional plots confirmed the distinct landscapes (**Fig. 4B-G**) and density profiles (**Fig. 4H-J**) of the 16S rRNA gene seqverses of the three domains of life. The seqverses revealed distinct multi-modal density distributions in the 2D coordinate space, as seen for the archaeal seqverse (**Fig. 4H**) or bacterial seqverse (**Fig. 4I**), around their respective domain centroids. The coordinates of the centroids for each domain were calculated as the median coordinate on each dimension using all sequences whose information at the phylum level was available (n= 331,105 cases). As expected the most distant sequences were those of *Pyrobaculum* (Archaea) and Phylum Foraminifera (Eukaryota), which contributed to the large spread of the archaeal and eukaryotic seqverses away from their centroids, respectively. The centroid (coordinates 0.017664, 0.094707, 0.046474) of the Archaea seqverse had the sequence of Thermoplasmata (identified as KP091008.1.1423 in the MSA) as closest sequence (distance of 0.0002192) (**Movie 3**; **Fig. 4E, H**). For bacteria, the closest to the centroid (coordinates x=0.010409, y=0.097673, z=0.051626) was the sequence of the uncultured Actinobacteria of the genus *Blastococcus* (identified as KC554984.1.1516) at a distance of 3.734e-05 (**Movie 4**; **Fig. 4F, I**). For Eukaryota, the closest to the centroid (coordinates 0.028267, 0.087945, 0.063611) belonged to the Animalia *Bilateria, Cnidaria, Ctenophora* (BCP; KY077283.1.1814) at a distance of 4.294182e-05 (**Movie 5**; **Fig. 4G, J**). The average distance of all sequences to their domain centroids was 0.00209 ± standard deviation 0.001867 for Archaea, 0.00395 ±0.002892 for Bacteria, and 0.00310± 0.004643 for Eukaryota, while the distances between domain centroids were on average 0.01808 ± 0.007626 (**Fig. 5A-B**). Therefore, the between-domain distances were on average 9.7, 4.6, and 5.80 times larger than the respective within-domain distances, underlying the sequence cohesiveness around domain centroids.

**Figure 5.**
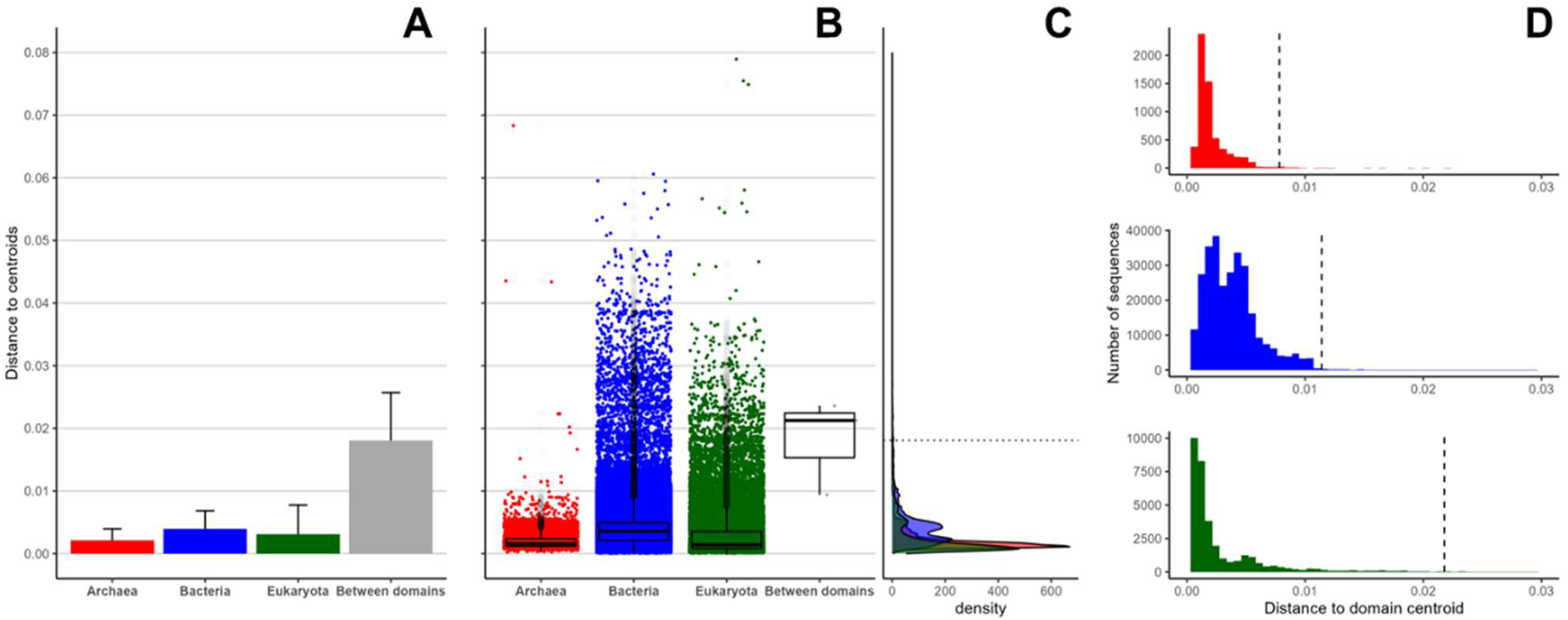
Between– and within-domain distances to centroids. Using the CAPASYDIS 3D distances of the NR99 dataset, **A**) the centroids of each domain seqverse were calculated and the within-domain distances to the centroids are depicted as barplot (means and standard deviation), as well as the average distance between the domain centroids (grey bar). Given the nature of the dataset, the numbers of sequences for each domain are unbalanced (with 6,031, 291,195, 34,437 points for Archaea, Bacteria, and Eukaryota, respectively), and with only three points corresponding to the distances between the domain centroids. **B**) A boxplot visualization of the same data with jittered points and their **C**) marginal density plots. The dotted horizontal line represents the mean distance between centroids. Seven points are not shown in panel B) because they are located between 0.08-0.13. **D**) Distribution of distances to the centroid for each domain (Archaea in red; Bacteria in blue; Eukaryota in green) with a dashed line representing the 99^th^ percentile for each distribution.

The distribution of distances to centroids for the domain Bacteria displayed a bimodal distribution, which was not evidenced for the other domains (**Fig. 5D**). To investigate this pattern, bacterial sequences were then divided into two groups, based on whether their distances to the centroid were smaller or larger than the observed distance cutoff of 0.0033, which was found as the mid distance between the two modes of the distribution (**Fig. S1**). Group 1 and group 2 included 134,978 (46.4%) and 155,744 (53.6%) bacterial sequences, respectively. To understand whether these groups were specific to particular taxonomic levels, detailed analysis at the Phylum level for each group was undertaken: A partitioning at the phylum level was observed for 22 out of 51 (43.1%) phyla. For those 22 phyla, >90% of their sequences were classified into one specific group, with group 1 with 21 cases and group 2 with 1 case corresponding to Pseudomonadota. The latter case corresponded to 6,545 (8.0%) and 75,748 (92.0%) sequences belonging to each group, respectively, suggesting a clear trend of association with group 2 for Pseudomonadota, i.e. the more diverging group of sequences to the domain centroid (**Table S1A**). Similarly, 41/105 (39.0%) of the bacterial Classes were associated with >90% of their sequences belonging to a specific group (group 1 with 37 cases and group 2 with 5 cases; **Table S1B**). Those patterns suggest that few bacterial Phyla and Classes may tend to systematically diverge from their domain centroid more than others. This observation is also supported by the 2D analyses (**Fig. 4F, I**), which indicated that the point density distribution of the bacterial domain was not uniformly distributed, as some denser areas were evidenced, much like an archipelago of small “islands” around a larger island where the domain centroid was located. In the 2D plots (**Fig. 4B-D**), the area of the Archaea seqverse was also 5.6 and 18.1 times smaller than that of the Bacteria and Eukaryota, respectively (area for Archaea: 0.00022, Bacteria: 0.00124, Eukaryota: 0.00401 asymdist^2^), suggesting a smaller diversity space for Archaea. Differences in domain shape were mostly driven by the presence of few “atypical” sequences, whose effects on the distributions were mostly seen for the Archaea (**Fig. 4E**) and Eukaryota (**Fig. 4G**) domains.

Atypical sequences in each domain were then arbitrarily defined as sequences whose distances to their domain centroid were greater than the respective 99% percentile (q99) of all the 3D distance values for this domain. The q99 values of 7.8 ×10^-03^, 11.4 e^-03^, and 21.8 e^-03^ were obtained for the domains Archaea, Bacteria, and Eukaryota, respectively, which mirror their observed spreads and shapes (**Fig. 5D**), and total 3D variance of 7.40 e-06, 2.36 e-05, and of 2.84e-05, respectively. At these q99 values, 61, 2,912 and 344 atypically distant sequences were observed for each domain, respectively. For the archaeal domain with 61 outliers, the phylum Thermoproteota was represented with classes Thermoproteia (n=39 sequences; 63.9%), and Bathyarchaeia (3 sequences; 4.9%), the phylum Aenigmarchaeota with the class “Deep Sea Euryarchaeotic Group” (DSEG) (8 sequences; 13.1%), and the phylum Thermoplasmatota with class Thermoplasmata (2 sequences; 3.3%). For the domain Bacteria, with 2,912 sequences with distances above q99, the top three phyla were phylum Bacillota with classes Clostridia (787 sequences; 27%), Bacilli (269 sequences; 9.3%), Negativicutes (104 sequences; 3.6%), Desulfotomaculia (101 sequences; 3.5%), and Desulfitobacteriia (65 sequences; 2.2%), phylum Pseudomonadota with classes Alphaproteobacteria (602 sequences; 20.7%), Gammaproteobacteria (46 sequences; 1.6%), and phylum Patescibacteria with classes Saccharimonadia (299 sequences; 10.3%), Parcubacteria (59 sequences; 2%) and Microgenomatia (57 sequences; 2%). For the domain Eukaryota with 345 outlier sequences, the identified phyla were Discoba with class Discicristata (143 sequences; 41.6%), phylum Amorphea with classes Obazoa (99 sequences; 28.8%) and Amoebozoa (18 sequences; 5.2%) and phylum Archaeplastida with classes Chloroplastida (45 sequences; 12.8%) and Rhodophyceae (26 sequences; 7.6%), and phylum SAR with class Rhizaria (13 sequences; 3.8%).

### **Seqverse of** *Enterobacteriaceae*

Because we chose *E. coli* as the first reference for the axis definition of the rRNA gene seqverse, it was essential to evaluate how effectively sequences from related species are represented in the resulting seqverse. We determined how sequence coordinates in the low-dimensional projection would align with various levels of taxonomic resolution. We thus considered the bacterial Family Enterobacteriaceae to which E. coli belongs, and kept the top 15 genera whose sequence abundances in the dataset consisted of at least 10 sequences. The clustering of the sequences by genus indicated that along the first two dimensions of the CAPASYDIS projection axes a continuum of diversity was observed (**Fig. 6**). Individual projection of each single genus on each axis (**Fig. S2**) clearly revealed that most Enterobacteriaceae genera, including Escherichia, Enterobacter, Klebsiella, Kluyvera, Salmonella, Shigella, of the Enterobacteriaceae displayed very peaky distributions on each projection axis, suggesting well-defined seqverses at the genus level. On the third axis, however, more overlap between the coordinates of different genera was observed. The lack of resolution observed for E. coli and relatives on the third axis could be expected, given that this axis was built using the most distant sequences to E. coli, which were the Eukaryotic (H. depressa) and the Archaeal *Pyrobaculum* sequences.

**Figure 6.**
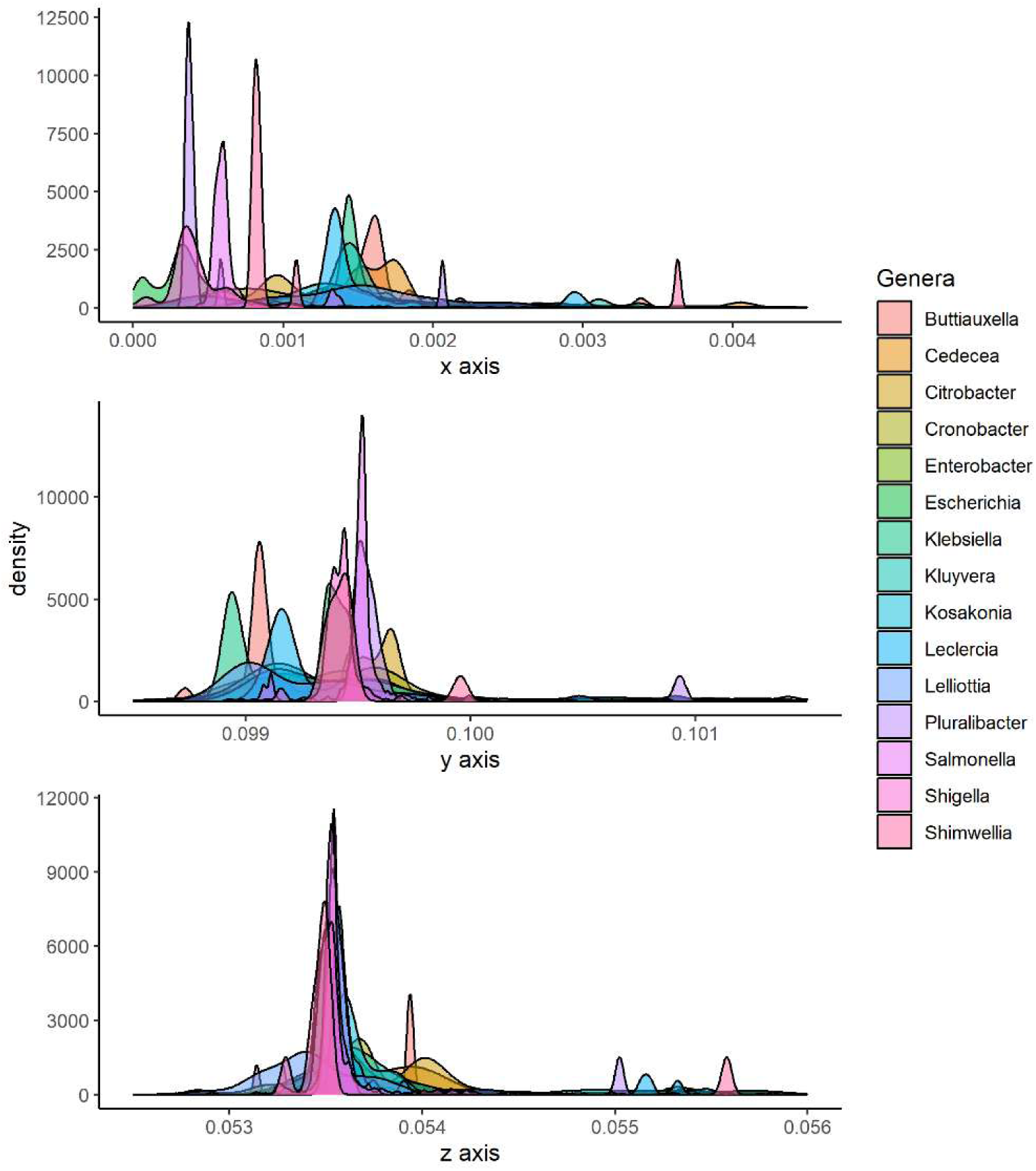
Density curves of sequence positions of the top 15 most abundant genera of the family Enterobacteriaceae onto individual CAPASYDIS axes.

### Seqverse of *E. coli*

When considering E. coli sequences, consisting of 948 of the 4,343 total Enterobacteriaceae sequences (**Fig. 7A**), a main “island” of diversity was found with two main peaks (**Fig. 7B**). To investigate this particular grouping, we focused on the subset of 200 E. coli sequences that differed by only one nucleotide difference from each other. The rationale to focus on this set was to evaluate how the smallest possible differences can be represented in the seqverse in comparison to other methods. Noticeably, the observed grouping of sequences also matched the two main clusters obtained with classical phylogenetic approaches (**Fig. 7C**), using classical dendrogram and unrooted tree representations (**Fig. 7D**). We further compared the number of base differences (Hamming distance) between all 948 E. coli sequences, and identified 655 pairs involved a single nucleotide difference, with a median of 15 and a mean of 22 nucleotide differences, and a maximum of 385 nucleotide differences (data not shown). Overall, many sequences could not be distinguished from each other with standard phylogenetic analyses, because they had the exact same base pair difference, even though these differences occurred at different positions along the aligned sequences. Taking the example of only one base difference between a sequence and the reference, the mismatched pair “TC” occurred in 134 sequence pairs (20.5%) and “CT” in 60 pairs (9.2%), which could not be distinguished from each other using classical Phylogenetics. With asymdist computations, all those single differences occurring at distinct sequence positions were translated into distinct coordinate values.

**Figure 7.**
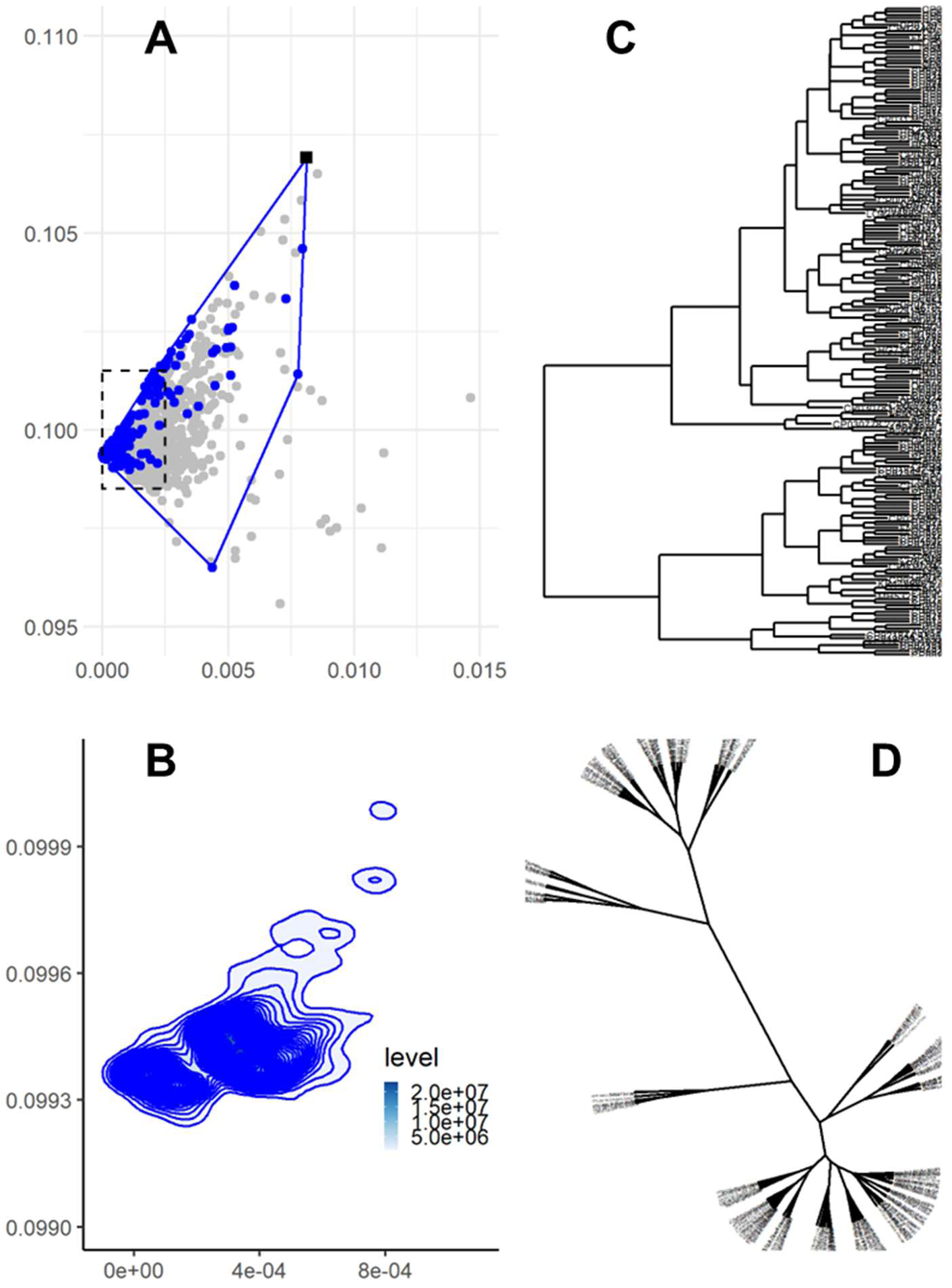
Visualization of *E. coli* seqverse within the Enterobacteriaceae. **A**) Within the Enterobacteriaceae (n = 4,408 sequences) colored in grey, the seqverse of E. coli sequences (n= 948) is indicated as blue points, as well as its corresponding convex hull. The black square represents the only sequence that was found at more than 99^th^ percentile from the centroid (AOQL01000076.469.2061). The dashed rectangle indicates the coordinates of the region shown in panel B. **B**) Contour plot (bins= 50) of the E. coli seqverse with density lines. **C**) Rectangular phylogram of the 200 E. coli sequences that only differed by one nucleotide change from each other. The ML tree was inferred from the alignment of 29,932 nucleotide sites with the TIM2+F+I model of substitution implemented in IQ-TREE (version 2.1.2). **D**) An unrooted version of the rectangular phylogram shown in panel C.

### Comparison of seqverse distances to phylogenetic distances

We further investigated whether the distances between the mapped points in the CAPASYDIS seqverses were related to sequence differences. We found that the number of nucleotide differences was significantly correlated with the distances between sequences in the 1D, 2D, and 3D CAPASYDIS seqverses, with Pearson correlation coefficients of 0.643, 0.640, and 0.640, respectively (all P<0.001; **Fig. S3**). Additionally, the maximum likelihood (ML) distances obtained with TIM2+F+I model in IQ-TREE (25) were compared to the distances obtained between the 1D, 2D and 3D seqverses of the 948 E. coli sequences consisting of 29,932 aligned nucleotide sites (analyses available in the GitHub repository). Of note, the distances estimated by ML-based models and asymdist are not directly comparable, as the former treat gaps (“-”) as unknown characters, essentially ignoring them during likelihood calculations and ignoring them for the site-likelihood calculation of a given tree. In the classical approach, a pair that involves a gap “-” (e.g. “A-”) is given a distance of 0.0000010 versus a distance of 0.0012879 (1,287 times larger) for the “AG” mismatch. This is in contrast to the calculations used in CAPASYDIS, where a specific weight is also given to cases where a gap is involved at a position (**Table 1**). Overall, the calculated ML distances fell into two groups, with a range of values from 0 to 0.003, and there were only 27 out of 655 (4.1%) distances which were numerically unique, illustrating the lack of discrimination already evidenced at the distance matrix level among closely related sequences. These 655 distances between the corresponding sequences in the seqverses consisted of 441 (67.3%), 516 (78.8%), and 585 (89.3%) unique distances (at the e-10 precision level) when considering the 1D, 2D or 3D seqverses, respectively. This means that, despite the compression of the information in a low-dimensional projection space, very small nucleotide differences and evolutionary distances between sequences are well represented in the rRNA seqverse.

### Comparison to alternative computational approaches

The CAPASYDIS approach was also compared to alternative computational methods that are often used to map at set of sequences in a reduced space (**Table 2**). This list of methods includes entropy-based, manifold learning and deep learning /Large Language-based approaches, and is intended to illustrate the most salient points introduced by each kind of approaches. Beyond classical phylogenetic approaches (see above), another bioinformatic solution could be to calculate a weighted Euclidean distance matrix between sequences based on their information content. For instance, by focusing on low-entropy columns (conserved regions) and penalizing mismatches there, one could thus weight “evolutionary constraints” in the MSA (26, 27). A mutation in a highly conserved active site would count for more than a mutation in a highly variable loop. Using Shannon entropy (28), sequences that deviate in conserved areas would thus stand out as outliers. One could apply a weight wj that is inversely proportional to the entropy value, to ensure that a mismatch in a low-entropy column results in large change in distance, whereas a mismatch in a “noisy” (high entropy) column would largely be ignored. The resulting sequence coordinates would be then further projected into an Euclidean space (like a “weighted” PCA). In that representation, the resulting coordinates would no longer just be unique identifiers (like in CAPASYDIS), but the axes actually would represent gradients of conservation (inversed entropy). Yet, this entropy-weighted approach would inherently create a dynamic system of coordinates because the addition of new sequences would alter the global probability distribution, and would imply recalculating column weights for the entire dataset. Adding or removing input sequences will also trigger each time a new rotation of the PCA axes that will adjust their extraction of the new total variance, effectively moving the “grid lines” of the projection map (29). Thus, the method based on weighted Euclidean distances of entropy information may provide interesting biological meaning, but at the cost of the stability of the representation. Additionally, phylogenetic and information-theoretic approaches incur high computational overhead. As the number of input sequences (N) grows, the cost of constructing distance matrices rises quadratically (O(N^2^)), while tree-building algorithms like Neighbor-Joining scale cubically (O(N^3^)) (30).

**Table 2.** Comparison of CAPASYDIS with possible alternative approaches to visualize biological sequences in a reduced space.

|  | CAPASYDIS | Classical<br>Phylogenetics | Entropy-based<br>approach | Manifold<br>Learning | Deep learning,<br>autoencoders,<br>Large Language<br>Models |
| --- | --- | --- | --- | --- | --- |
| Primary goal | Spatial<br>indexing and<br>mapping | Evolutionary<br>Inference | Evolutionary<br>outlier detection | Local clustering<br>and transition | Functional<br>classification;<br>feature extraction<br>and fitness |
| Input type | Aligned<br>sequences | Aligned<br>sequences | Aligned<br>sequences | Aligned<br>sequences | Unaligned<br>sequences |
| Mathematical<br>basis | Asymmetric<br>distances | Symmetric<br>genetic<br>distances | Information<br>theory (Shannon<br>entropy) per<br>column to derive<br>weights | Symmetric<br>genetic<br>distances;<br>Laplacian<br>eigenmaps | Non-linear Neural<br>mapping |
| Representation | Cartesian<br>projection | Phylogenetic<br>trees;<br>dendrograms | Classical<br>dimensional<br>reduction (e.g.<br>weighted PCA) | Projection of<br>the first<br>eigenvectors;<br>cluster analysis<br>(e.g. GMM) | Latent space<br>trajectories;<br>cluster analysis<br>(UMAP, t-SNE) |
| Information<br>type | Geometric/<br>structural | Ancestral/<br>historical | Conservation/<br>unexpected<br>signal | Probabilistic<br>density | Biological<br>"Grammar" |
| Coordinate<br>stability | Absolute:<br>Fixed<br>addresses | Relative to<br>input and tree<br>topology | Relative to input | Stochastic:<br>Relative to<br>neighborhood | Deterministic:<br>Model-fixed logic |
| Scalability | High (linear /<br>$O(N)$) | Low to<br>moderate:<br>computational<br>cost of<br>repeating<br>calculations<br>upon new<br>entries<br>($O(N^2)$,<br>$O(N^3)$) | Low to moderate:<br>computational<br>cost of repeating<br>calculations upon<br>new entries<br>($O(N^2)$) | Low to<br>moderate:<br>(graph<br>construction);<br>($O(N^2)$) | Moderate (training<br>cost) |

Abbreviations: **GMM**, Gaussian Mixture Models; **NJ**, Neighbor Joining; **PCA**, Principal Component Analysis; **t-SNE**, t-distributed Stochastic Neighbor Embedding; **UMAP**, Uniform Manifold Approximation Projection.

Another geometric possibility would be to embed the aligned sequences from the MSA into a vector space using, for instance, Laplacian eigenmaps (manifold folding), followed by Gaussian Mixture Models (GMM) to cluster the sequences probabilistically (**Table 2**), as illustrated for instance by Bruneau et al. (31). The workflow begins by computing an all-versus-all distance matrix based on the aligned sequences in the MSA (i.e. symmetric distances), and using Laplacian Eigenmaps, local geometry is prioritized by projecting sequences into a space that keeps close biological “neighbors” together, creating a high-fidelity manifold (32). The low-dimensional environment then allows GMM to calculate membership probabilities, providing a statistical view on how sequences may transition between delineated groups. A major difference with CAPASYSIS is the limited scalability of the “Laplacian Eigenmaps/GMM” approach, which does not avoid the “curse of dimensionality”, as all-versus-all distance matrix also needs to be computed. In addition, commonly used clustering methods such as GMM, t-SNE or UMAP are stochastic (different local optimal may be found) and may produce different sets of sequence coordinates each time the algorithms are run on the same dataset depending on the “neighborhood” of data points and on the chosen initialization parameters (33–35).

A non-linear/generative alternative would be to apply autoencoders to the data present in a MSA (**Table 2**). Aforementioned approaches and CAPASYDIS itself all require a high-quality MSA as input, otherwise noisy sequences are expected to distort the appearance of the rendered sequence spaces, and the methods cannot mix sequences of different lengths (e.g., short and long reads) easily. On the contrary, deep learning/Large Language Models (LLMs) approaches can handle “unaligned” sequences”, as LLMs like DNABERT (36) or Nucleotide Transformer (37), learn directly from raw tokens (k-mers). Deep learning is a broad technique that uses multi-layered artificial neural networks to learn patterns from data. A specific type of deep learning network design, autoencoders, is often used to compress data and then reconstruct it, with useful applications for dimensionality reduction or noise removal (38, 39). LLM algorithms, which are built using deep learning architectures (specifically transformers), do not just measure distance between the sequences. They learn the syntax of the sequences (e.g., promoter regions, motifs), and thus produce a complex, non-linear mapping function (40, 41). The study by Yeung et al. provides an example that leverages deep learning to represent sequences with various levels of complexity and similarity (42). Pre-trained Protein Language Models (PLMs) were used to convert protein sequences directly into high-dimensional numerical vectors (embeddings), without needing explicit alignment, thereby capturing latent structural and functional grammar from millions of sequences. These vectors were then analyzed using Neighbor-Joining (NJ) trees or dimensionality reduction tools like Uniform Manifold Approximation and Projection (UMAP). Once trained and frozen, a neural network functions as a deterministic map, assigning a specific input sequence to the exact same latent vector every time (40). Since these coordinates are determined by fixed model weights rather than by pairwise comparisons with other data points, inference is highly efficient and scales linearly with sequence length. This efficiency yet relies on the network having ‘encoded’ evolutionary rules (e.g. deep biological similarities, shared motifs) during its computationally intensive training phase. However, this parametric approach introduces a rigidity that CAPASYDIS avoids: Because an autoencoder is bound by the logic of its training set, if trained only on Bacteria, it may attempt to force an archaeal sequence into a ‘bacterial’ representation. This typically would result in an out-of-distribution error, where the model fails because the input is fundamentally different from the data it studied during training (43). Instead, CAPASYDIS processes sequences based on their intrinsic information, without referring to a training set. If a novel organism (e.g., an Archaeon) is added to a Bacterial dataset, it simply projects it to a distant and specific coordinate in the seqverse. It is important to note that, although generating embeddings via PLMs is scalable (linear per sequence), the visualization step still faces bottlenecks. Indeed, calculating NJ trees or UMAP plots for massive datasets are computationally expensive (O(N^2^) or O(N^3^)). Yeung et al. (42) explicitly noted that “tree-based methods may not scale as well to larger datasets” compared to dimensionality reduction. Furthermore, these visualizations (dendrograms) are topologically informative but visually “non-unique”, as branches or group of points can be rotated without changing the information. This issue is explicitly addressed with the CAPASYSIS method.

Overall, the comparative analysis to other computational approaches (**Table 2**) demonstrates that CAPASYDIS fills a critical niche for the linear (O(N)) spatial indexing and visualization of massive sequence datasets. While alternative computational methods may capture richer biological context, they suffer from the ‘curse of dimensionality’ and often fail to distinguish positional variants, sequences that differ only by the location of their mutations. By ensuring that every unique sequence variation yields a mathematically distinct coordinate, CAPASYDIS establishes an absolute, fixed reference frame. This guarantees stable positioning that remains independent of future dataset updates, solving the scalability issues inherent to dynamic mapping approaches.

## Discussion

CAPASYDIS is a computational framework designed to map aligned sequences from a multiple sequence alignment into a low-dimensional space where every distinct sequence receives a fixed, unique, non-overlapping coordinate. This is achieved through the concept of “asymmetric” distances, which applies specific weighting to prevent mathematical collisions, a scenario where different sequences might otherwise result in identical coordinates. CAPASYDIS chooses main axes of reference a priori to avoid the “curse of dimensionality”, unlike traditional dimensionality reduction methods like PCA, t-SNE, and UMAP (8) or tree-like representation (e.g. classical Phylogenetics; (30, 44)) that rely on all-versus-all matrix computation, or deep learning models that require massive training datasets (45, 46).

The mathematical foundation of the obtained coordinates relies on specific *delta* values that map events uniquely and to the mathematical properties of the associated radicands (e.g., 1+delta, 2+delta, 3+delta,…). The magnitude of the *delta* values is critical, as it governs how biological information is weighted in the *asymdist* computation. We set *delta* values that are 10 to 1,000 times smaller than the base index value of 1. This specific scaling was necessary to strike a balance between “uniqueness,” ensuring distinct mutation types are distinguished (**Table 1**), and “proportionality,” ensuring that the numerical contribution of a change at a position remains minute compared to the score obtained when bases are identical. The choice of the function involved in the calculation of *asymdist* was found empirically by trials and errors, but other mathematical functions could be envisioned, given that they map each unique event uniquely into a resulting coordinate space, and ideally provide a distance metric based on biological properties. The reverse function, i.e. going from a random point on the seqverse back to a sequence of bases, is less trivial, because points are not randomly located in a seqverse and the identification of a point would need to be made at the same precision as the map, e.g. e-10 in our study.

The second foundational consideration in the CAPASYDIS framework is “coordinate uniqueness”, i.e. whether the combinatorial density of the sequence projection space (“seqverse”) leads to analytical collisions, where biologically distinct mutation patterns produce identical spatial coordinates. We thoroughly address this structural boundary through a combination of algebraic field theory and multi-dimensional geometry testing (section “Algebraic independence and exhaustive matrix audit of score uniqueness”; Supplementary Material): In summary, for two distinct mutational patterns to yield identical scores on a single analytical axis, their respective sums of radicals would need to be rationally dependent over the field of rational numbers ℚ. Under Besicovitch’s Theorem (47), radicals of integers are strictly linearly independent over ℚ if their radicands (e.g., 1+delta, 2+delta, 3+delta,…) share no common square-free factors. To test this systematically under our dynamic configuration of *delta* values (**Table 1**), we executed an exhaustive global matrix audit of all 100 baseline position (index)-*delta* building blocks. This audit revealed exactly seven instances of structural rational dependence in a single dimension, rendering the 1D projection theoretically vulnerable to overlapping coordinates, given the suggested *delta* values in Table 1. However, CAPASYDIS resolves these linear dependencies by utilizing independent biological reference points to generate multi-dimensional (2D or 3D) coordinate vectors. For a spatial collision to occur, two distinct mutational states must collide across all orthogonal reference axes simultaneously. Scaling our global audit to a 2D framework confirmed that these 1D rational dependencies completely disperse into distinct spatial signatures, resulting in zero spatial coordinate collisions across the entire combinatorial continuum. This structural independence was also empirically validated by a brute-force analysis of the fully mutated M10bases dataset consisting of 1,048,576 distinct sequences (Supplementary Material), which yielded 100% numerically unique coordinates in 2D and 3D spaces. Furthermore, we verified that the analytical separation “gap” between unique multi-axis scores remains safely within the limits of machine epsilon for standard 64-bit floating-point precision (approx. 1.11 x 10^-16^). This is a well-characterized limit in computational geometry known as the “Sum of Square Roots” problem (48, 49). Collectively, these algebraic constraints and multi-axis geometries ensure that CAPASYDIS provides an ultra-dense, yet strictly non-overlapping and computationally resolvable representation of massive sequence spaces.

To ideally represent the resulting seqverses, we thus recommend using a low number of dimensions, specifically three for the rRNA case study, to allow for easy user interaction and visualization. The selection of reference sequences directly dictates the mapping results. For example, our study arbitrarily selected the E. coli as the initial point of origin of the rRNA seqverses, as it is one of the most extensively studied organism on Earth (50). From this starting point, we iteratively calculated the most divergent sequences to establish three main axes through triangulation, which enabled the spatial representation of very divergent sequences originating from the three domains of life in the large rRNA dataset. This system offers significant flexibility: Researchers can use the most distant references to create global diversity maps, as done for the NR99 dataset, or restrict references to a narrow variability space for targeted micro-evolution studies. Adopting a standardized reference system, similar to orienting maps with “North at the top,” would allow different research groups to integrate findings consistently across experiments and studies.

A primary advantage of this new framework is its scalability, as new sequences can be placed efficiently into an existing sequence space without the need to re-calculate the entire set of coordinates for already mapped sequences. However, this stability is strictly bound to the quality and structure of the underlying MSA, as any change in MSA positions would result in a new set of coordinates for the whole dataset. MSA reconstruction is known to scale poorly (O(N^2^) or O(N^3^)) and to struggle with highly divergent sequences (51, 52). In our study, we addressed this main limitation by using a large and stable reference MSA, spanning approximately 50,000 positions, capable of accommodating highly variable regions without altering the global MSA structure (14). To add and align new sequences to these extensive MSA without altering the global template, tools like the SILVA Incremental Aligner (SINA), which utilizes k-mer distance search and Partial Order Alignment (POA), have proven to be a versatile and robust tool (53). Rather than relying on a global template built from all references, SINA dynamically selects a small, fixed number of optimal reference sequences to construct a local alignment template. This approach significantly improves alignment quality, minimizes unaligned bases, and allows the method to scale efficiently to very large reference MSA.

Biologically, the maps generated by CAPASYDIS using the NR99 dataset correlate significantly with known taxonomic relationships. In testing, the seqverses successfully clustered sequences into the three known domains of life, confirming that Archaea are distant from the bacterial reference *E. coli*, with Eukaryotes being the most distant group. This reflects general knowledge about domain organization in the Tree of life (54–56). The three domain-specific seqverses differed in shapes, sizes and patterns of variation around their centroids. The detailed analyses for the bacterial phylum, order or family levels revealed that closely related genera display a “continuum” in their seqverses, as illustrated by the case of Enterobacteriaceae. For specific bacterial taxa, we also identified diversity patterns associated with more systematic divergence from the rest of the sequences of the domain Bacteria, suggesting the existence of either diversification patterns specific to those taxa or some difficult placement of rRNA sequences for these organisms. Overall, not only were the sequence coordinates in the seqverses taxonomically meaningful, but also the distances between points in the seqverses preserved meaningful information related to sequence differences or phylogenetic distances. Investigating the underlying causes of the identified diversification patterns lies beyond the scope of this study, but would be an interesting avenue for future research. While closely related bacterial genera displayed a “continuum” in the seqverse, we also noted that some regions remain unpopulated due to functional constraints. Specifically, the complex secondary and tertiary folding of rDNA molecules prevents mutations that would disrupt these essential structures (57). As the Tree of Life continues to be a central organizing principle in biology, our visualization of the existing sequence universe should grow in a scalable manner, i.e. in a manner that can cope with the ever-increasing amount of data generated by sequencing. Despite meaningful biological correlations, we emphasize that the asymmetric distance values are not intended to serve as strict phylogenetic or evolutionary distances. Moreover, the resulting seqverses based on either the forward or reverse sequence directions of the same dataset are simply not “mirror” images of each other, and cannot be derived by simply rotating one seqverse coordinate to produce the other representation (Supplementary Material).

Importantly, the asymmetric CAPASYDIS positioning aligns with fundamental molecular biology: Because template-directed enzymes (such as polymerases and ribosomes) process nucleic acids and proteins with strict chemical directionality (e.g., 5’→3’ or N– → C-terminus), mutations at distinct positions carry asymmetric biological weight depending on their linear context, a feature natively captured by the CAPASYDIS coordinate architecture. Although the forward-based and reverse-based sequence directions produce distinct seqverses, the latter are highly correlated to each other, with linear correlation coefficients ranging from 0.60 (2D, 3D) to 0.78 (1D) in the M10bases dataset. This behavior is expected given the asymmetric nature of the CAPASYDIS distance calculations, which incorporates sequence positional information and hence provides a complementary view of a given multi-sequence dataset. Unlike classical phylogenetics, however, CAPASYDIS does not distinguish between evolutionary timescales, such as recent versus ancient divergence, and does not compare all sequences with each other. Consequently, the resulting groups may not align perfectly with traditional phylogenetic reconstructions of species or Operational Taxonomic Units (OTUs). In addition, the resulting mapping of rRNA sequence differences in the NR99 seqverses may be disconnected from biological phenomena like horizontal gene transfer across genomes including the 16S rRNA gene itself (58), bacterial lifestyle or pangenome fluidity (59), and uneven evolutionary rates across different regions of the rRNA genes (60, 61).

In conclusion, CAPASYDIS helps define the boundaries of known molecular diversity, or “terra cognita”, and offers several avenues for future applications. These include overlaying metadata such as predicted functions or phenotypes onto the seqverses, or using the system for diagnostics to rapidly score long-read sequences for pathogen discovery. Although developed on DNA sequences, future applications to RNA or amino acid sequences could also be foreseen. Finally, because the approach allows for precisely positioning any sequences in a fixed coordinate system, the location of artificially generated sequences in a seqverse could be predicted in order to identify novel regions of interest.

## Acknowledgements

The idea of an entire galaxy existing within a marble in the 1997 film “Men in Black” served as a key inspiration for this work on dimensionality reduction, consistent mapping, and interactive visualization.

