## Supplementary Material for "Unveiling the Terra Cognita of Sequence Spaces using Cartesian Projection of Asymmetric Distances"

Alban Ramette[1,2]

**Affiliations**
[1] Institute for Infectious Diseases, University of Bern, Bern, Switzerland
[2] Multidisciplinary Center for Infectious Diseases, University of Bern, Bern, Switzerland

#### Figure S1

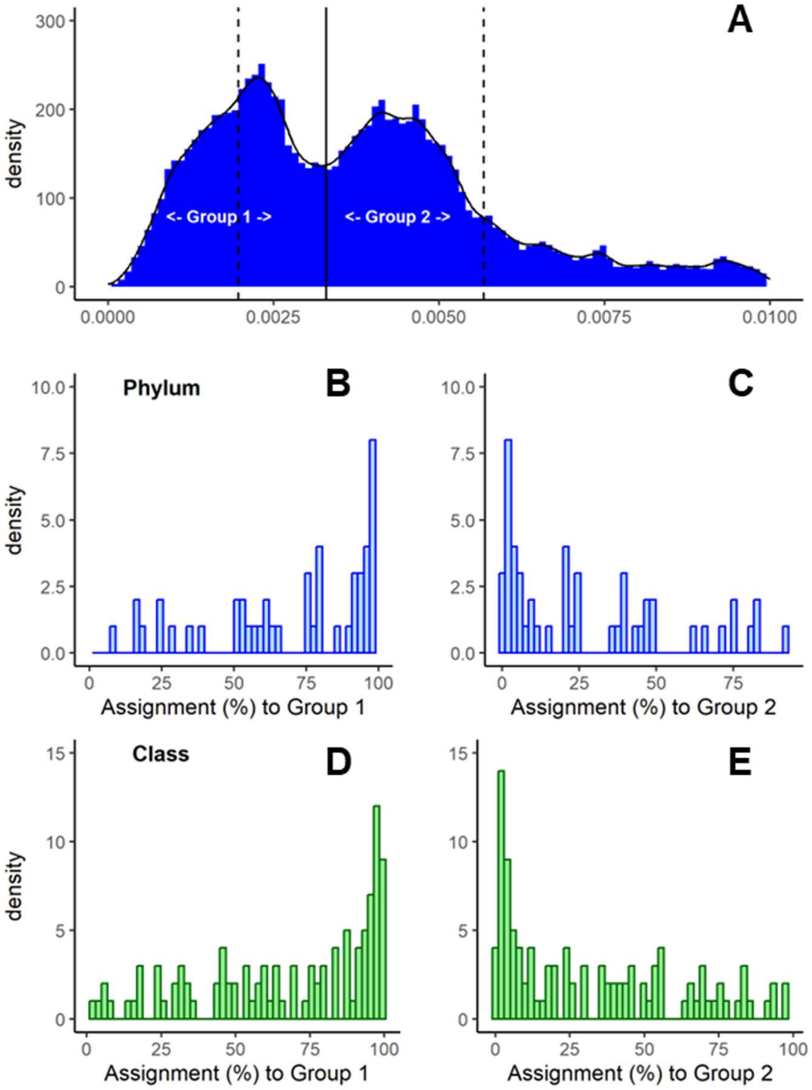

**Figure S1**. **Taxonomic analyses of the two modes observed in the distribution of 3D distances to the bacterial centroid in the bacterial seqverse. A**) The group definition is based on a cutoff of 0.0033 (thick black line), defined as the mid-point between the two observed peaks. Dotted lines are the means for each group of 0.001972, and 0.005671, respectively. **B-C**) Density distributions of the percentage assignment of bacterial sequences classified at the Phylum level for each group, respectively. **D-E**) Density distributions of the percentage assignment of bacterial sequences classified at the Class level for each group, respectively. Only levels with at least 100 sequences were considered here.

#### Figure S2

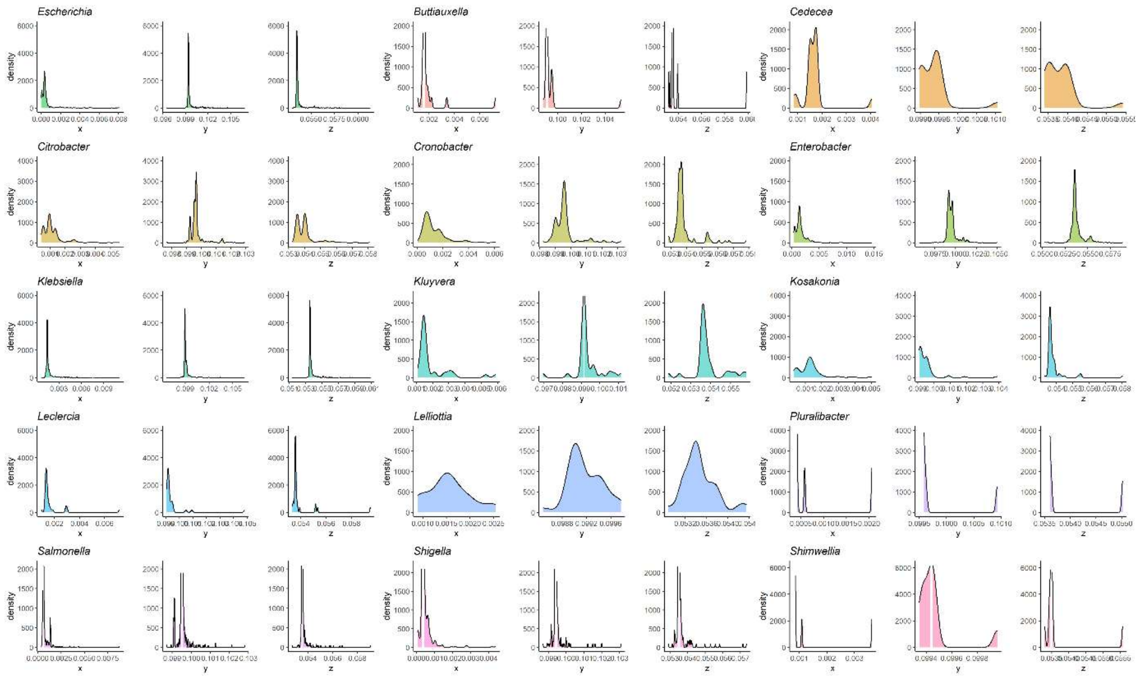

**Figure S2. Marginal densities of the coordinate distribution of sequences belonging to genera of the Enterobacteriaceae family for each CAPASYDIS axis.** Only the top 15 most abundant genera are shown. *E. coli* is shown as the first genus, and then the other genera are shown in alphabetical order.

#### Figure S3

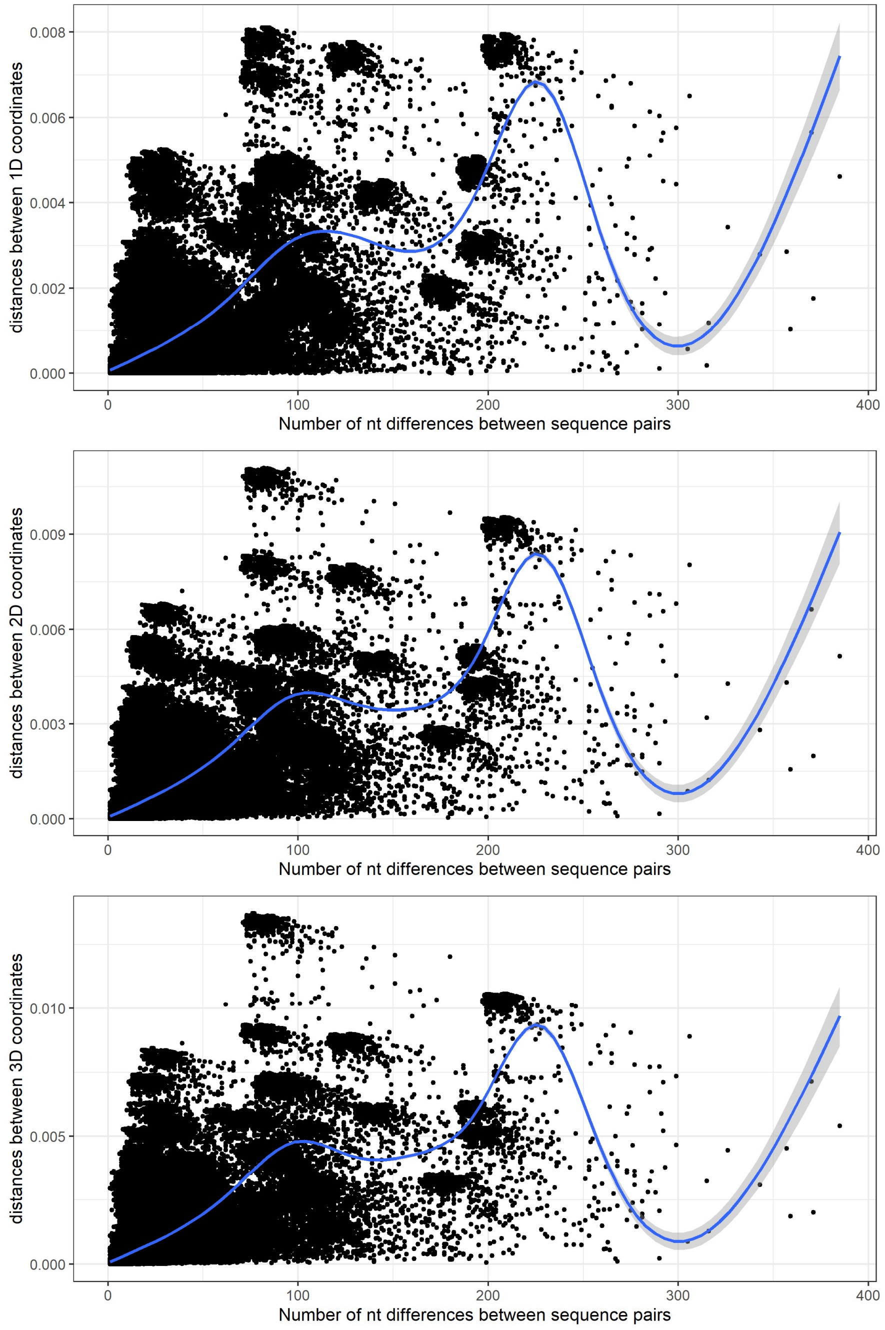

**Figure S3. Comparison of the distances between the sequence points in the (1D, 2D, 3D) CAPASYDIS seqverses (y axis) with the number of nucleotide differences between the same sequences (x axis).** The blue line depicts a non-linear smoothing curve (generalized additive model). The point density for pairs displaying more than 200 nucleotide differences is strongly reduced, which explains the drop in loess curve and larger confidence estimates after 250 nucleotide differences.

#### Table S1

##### A) Phyla present in each group defined in Figure S1

The phyla with more htan 90% sequences present in a specific group are display in bold font. Only phyla with more than 100 sequences are displayed here.

| | P1 | P1pc | P2 | P2pc | Total |
|---|---|---|---|---|---|
| | Group 1 | | Group 2 | | |
| Phylum | Count | Pc | Count | Pc | Total |
| **Sva0485** | **293** | **100.0** | **0** | **0.0** | **293** |
| **Ignavibacteriota** | **225** | **99.1** | **2** | **0.9** | **227** |
| **RCP2-54** | **108** | **99.1** | **1** | **0.9** | **109** |
| **NB1-j** | **406** | **98.8** | **5** | **1.2** | **411** |
| **Methylomirabilota** | **289** | **98.6** | **4** | **1.4** | **293** |
| **Thermodesulfobacteriota** | **5890** | **98.5** | **90** | **1.5** | **5980** |
| **Calditrichota** | **197** | **98.5** | **3** | **1.5** | **200** |
| **Dependentiae** | **473** | **98.3** | **8** | **1.7** | **481** |
| **Entotheonellaeota** | **113** | **98.3** | **2** | **1.7** | **115** |
| **Balneolota** | **306** | **98.1** | **6** | **1.9** | **312** |
| **Modulibacteria** | **150** | **98.0** | **3** | **2.0** | **153** |
| **MBNT15** | **121** | **96.8** | **4** | **3.2** | **125** |
| **Marinimicrobia** | **568** | **96.6** | **20** | **3.4** | **588** |
| **Rhodothermota** | **1624** | **96.2** | **65** | **3.8** | **1689** |
| **Acidobacteriota** | **9155** | **95.4** | **445** | **4.6** | **9600** |
| **Zixibacteria** | **161** | **94.2** | **10** | **5.8** | **171** |
| **Latescibacterota** | **377** | **94.0** | **24** | **6.0** | **401** |
| **Dadabacteria** | **138** | **93.9** | **9** | **6.1** | **147** |
| **Nitrospinota** | **254** | **92.0** | **22** | **8.0** | **276** |
| **Sumerlaeota** | **227** | **91.5** | **21** | **8.5** | **248** |
| **SAR324** | **478** | **91.0** | **47** | **9.0** | **525** |
| Hydrogenedentes | 182 | 89.2 | 22 | 10.8 | 204 |
| Actinomycetota | 23123 | 85.4 | 3964 | 14.6 | 27087 |
| Cloacimonadota | 128 | 80.0 | 32 | 20.0 | 160 |
| Spirochaetota | 2213 | 79.9 | 556 | 20.1 | 2769 |
| Bdellovibrionota | 1082 | 79.9 | 273 | 20.1 | 1355 |
| Nitrospirota | 781 | 79.6 | 200 | 20.4 | 981 |
| Campylobacterota | 2331 | 77.2 | 687 | 22.8 | 3018 |
| Verrucomicrobiota | 3646 | 76.4 | 1124 | 23.6 | 4770 |
| Synergistota | 530 | 75.2 | 175 | 24.8 | 705 |
| Elusimicrobiota | 237 | 75.0 | 79 | 25.0 | 316 |
| Fusobacteriota | 883 | 64.3 | 491 | 35.7 | 1374 |
| Chlamydiota | 154 | 63.1 | 90 | 36.9 | 244 |
| Planctomycetota | 3893 | 60.3 | 2559 | 39.7 | 6452 |
| Myxococcota | 1410 | 60.3 | 927 | 39.7 | 2337 |
| Thermotogota | 124 | 59.9 | 83 | 40.1 | 207 |
| Gemmatimonadota | 1017 | 57.7 | 745 | 42.3 | 1762 |
| Abditibacteriota | 79 | 54.1 | 67 | 45.9 | 146 |
| Bacteroidota | 17928 | 53.3 | 15712 | 46.7 | 33640 |
| Fibrobacterota | 243 | 52.3 | 222 | 47.7 | 465 |
| Chloroflexota | 3472 | 51.3 | 3297 | 48.7 | 6769 |
| Bacillota | 39288 | 51.1 | 37579 | 48.9 | 76867 |
| Armatimonadota | 219 | 38.4 | 352 | 61.6 | 571 |
| Patescibacteria | 1182 | 33.8 | 2314 | 66.2 | 3496 |
| Halanaerobiaeota | 58 | 29.4 | 139 | 70.6 | 197 |
| Deinococcota | 162 | 24.4 | 501 | 75.6 | 663 |
| Aquificota | 42 | 24.0 | 133 | 76.0 | 175 |
| Cyanobacteriota | 1417 | 18.7 | 6143 | 81.3 | 7560 |
| Acetothermia | 23 | 17.2 | 111 | 82.8 | 134 |
| Atribacterota | 53 | 16.8 | 262 | 83.2 | 315 |
| **Pseudomonadota** | **6545** | **8.0** | **75748** | **92.0** | **82293** |

#### B) Classes present in each group defined in Figure S1

| | Group 1 | | Group 2 | | |
|---|---|---|---|---|---|
| Class | Count | Pc | Count | Pc | Total |
| **Desulfobacteria** | 1847 | 99.6 | 7 | 0.4 | 1854 |
| **Syntrophia** | 350 | 99.4 | 2 | 0.6 | 352 |
| **BD2-11** | 328 | 99.4 | 2 | 0.6 | 330 |
| **Ignavibacteria** | 225 | 99.1 | 2 | 0.9 | 227 |
| **Desulfobulbia** | 1141 | 99.0 | 11 | 1.0 | 1152 |
| **Thermoleophilia** | 1394 | 98.8 | 17 | 1.2 | 1411 |
| **S0134** | 161 | 98.8 | 2 | 1.2 | 163 |
| **Methylomirabilia** | 289 | 98.6 | 4 | 1.4 | 293 |
| **Calditrichia** | 197 | 98.5 | 3 | 1.5 | 200 |
| **Babeliae** | 473 | 98.3 | 8 | 1.7 | 481 |
| **Entotheonellia** | 113 | 98.3 | 2 | 1.7 | 115 |
| **Balneolia** | 306 | 98.1 | 6 | 1.9 | 312 |
| **Moduliflexia** | 150 | 98.0 | 3 | 2.0 | 153 |
| **Syntrophobacteria** | 176 | 97.8 | 4 | 2.2 | 180 |
| **Desulfuromonadia** | 1091 | 97.6 | 27 | 2.4 | 1118 |
| **bacteriap25** | 282 | 97.6 | 7 | 2.4 | 289 |
| **Acidobacteriae** | 3095 | 97.2 | 88 | 2.8 | 3183 |
| **Desulfovibrionia** | 805 | 97.1 | 24 | 2.9 | 829 |
| **Thermoanaerobaculia** | 460 | 97.0 | 14 | 3.0 | 474 |
| **Blastocatellia** | 1452 | 96.9 | 47 | 3.1 | 1499 |
| **Vicinamibacteria** | 2536 | 96.5 | 92 | 3.5 | 2628 |
| **Rhodothermia** | 1624 | 96.2 | 65 | 3.8 | 1689 |
| **Brocadiae** | 113 | 95.8 | 5 | 4.2 | 118 |
| **Aminicenantia** | 314 | 95.7 | 14 | 4.3 | 328 |
| **Nitrospinia** | 201 | 95.7 | 9 | 4.3 | 210 |
| **Syntrophorhabdia** | 123 | 95.3 | 6 | 4.7 | 129 |
| **Latescibacteria** | 140 | 95.2 | 7 | 4.8 | 147 |
| **Leptospirillia** | 123 | 94.6 | 7 | 5.4 | 130 |
| **Bacteriovoracia** | 272 | 94.4 | 16 | 5.6 | 288 |
| **count_1** | 1066 | 94.1 | 67 | 5.9 | 1133 |
| **Dadabacteriia** | 138 | 93.9 | 9 | 6.1 | 147 |
| **Coriobacteriia** | 855 | 93.5 | 59 | 6.5 | 914 |
| **Pla4** | 120 | 93.0 | 9 | 7.0 | 129 |
| **Oligoflexia** | 481 | 92.0 | 42 | 8.0 | 523 |
| **Acidimicrobiia** | 2030 | 91.6 | 187 | 8.4 | 2217 |
| **Sumerlaeia** | 227 | 91.5 | 21 | 8.5 | 248 |
| **Rubrobacteria** | 160 | 90.9 | 16 | 9.1 | 176 |
| Hydrogenedentia | 182 | 89.2 | 22 | 10.8 | 204 |
| Nitrospiria | 382 | 87.8 | 53 | 12.2 | 435 |
| NA | 3361 | 87.6 | 477 | 12.4 | 3838 |
| Verrucomicrobiia | 2847 | 87.6 | 404 | 12.4 | 3251 |
| Endomicrobiia | 101 | 87.1 | 15 | 12.9 | 116 |
| MB-A2-108 | 135 | 86.5 | 21 | 13.5 | 156 |
| Actinobacteria | 18477 | 83.5 | 3647 | 16.5 | 22124 |
| Negativicutes | 2195 | 83.1 | 445 | 16.9 | 2640 |
| Spirochaetia | 1989 | 82.9 | 410 | 17.1 | 2399 |
| KD4-96 | 106 | 82.8 | 22 | 17.2 | 128 |
| Dethiobacteria | 81 | 80.2 | 20 | 19.8 | 101 |
| Cloacimonadia | 128 | 80.0 | 32 | 20.0 | 160 |
| Chitinivibrionia | 189 | 79.7 | 48 | 20.3 | 237 |
| Campylobacteria | 2306 | 77.1 | 685 | 22.9 | 2991 |
| D8A-2 | 105 | 76.6 | 32 | 23.4 | 137 |
| vadinHA49 | 173 | 75.5 | 56 | 24.5 | 229 |
| Synergistia | 530 | 75.2 | 175 | 24.8 | 705 |
| Desulfitobacteriia | 296 | 75.1 | 98 | 24.9 | 394 |
| Bacilli | 22902 | 73.4 | 8304 | 26.6 | 31206 |
| Holophagae | 371 | 70.5 | 155 | 29.5 | 526 |
| Lentisphaeria | 247 | 69.8 | 107 | 30.2 | 354 |
| Syntrophomonadia | 125 | 69.8 | 54 | 30.2 | 179 |

| | | | | | |
|---|---|---|---|---|---|
| Desulfotomaculia | 328 | 64.8 | 178 | 35.2 | 506 |
| Kiritimatiellia | 479 | 64.4 | 265 | 35.6 | 744 |
| Fusobacteriia | 883 | 64.3 | 491 | 35.7 | 1374 |
| Chlamydiia | 154 | 63.1 | 90 | 36.9 | 244 |
| Planctomycetes | 2214 | 61.7 | 1377 | 38.3 | 3591 |
| Bdellovibrionia | 329 | 60.5 | 215 | 39.5 | 544 |
| Thermotogae | 124 | 59.9 | 83 | 40.1 | 207 |
| Thermodesulfovibrionia | 196 | 58.9 | 137 | 41.1 | 333 |
| Polyangiia | 897 | 58.4 | 638 | 41.6 | 1535 |
| Vampirivibrionia | 218 | 57.2 | 163 | 42.8 | 381 |
| OM190 | 185 | 55.7 | 147 | 44.3 | 332 |
| Anaerolineae | 2164 | 54.1 | 1837 | 45.9 | 4001 |
| Abditibacteriia | 79 | 54.1 | 67 | 45.9 | 146 |
| Bacteroidia | 17928 | 53.3 | 15712 | 46.7 | 33640 |
| Phycisphaerae | 925 | 49.6 | 939 | 50.4 | 1864 |
| Dehalococcoidia | 609 | 49.6 | 620 | 50.4 | 1229 |
| Longimicrobiia | 91 | 48.7 | 96 | 51.3 | 187 |
| Gracilibacteria | 277 | 47.0 | 312 | 53.0 | 589 |
| Microgenomatia | 310 | 46.7 | 354 | 53.3 | 664 |
| Armatimonadia | 57 | 46.0 | 67 | 54.0 | 124 |
| Myxococcia | 231 | 45.0 | 282 | 55.0 | 513 |
| Ktedonobacteria | 168 | 45.0 | 205 | 55.0 | 373 |
| Leptospirae | 91 | 44.6 | 113 | 55.4 | 204 |
| Thermoanaerobacteria | 158 | 43.4 | 206 | 56.6 | 364 |
| ABY1 | 125 | 36.7 | 216 | 63.3 | 341 |
| Gemmatimonadia | 331 | 34.0 | 643 | 66.0 | 974 |
| TK10 | 50 | 33.8 | 98 | 66.2 | 148 |
| Saccharimonadia | 301 | 32.1 | 637 | 67.9 | 938 |
| Clostridia | 12592 | 31.3 | 27693 | 68.7 | 40285 |
| Dojkabacteria | 36 | 31.3 | 79 | 68.7 | 115 |
| Limnochordia | 111 | 30.7 | 251 | 69.3 | 362 |
| Halanaerobiia | 58 | 29.4 | 139 | 70.6 | 197 |
| Aquificia | 42 | 26.8 | 115 | 73.2 | 157 |
| Deinococci | 162 | 24.4 | 501 | 75.6 | 663 |
| Fibrobacteria | 54 | 23.7 | 174 | 76.3 | 228 |
| Chloroflexia | 125 | 23.1 | 417 | 76.9 | 542 |
| Alphaproteobacteria | 4552 | 17.7 | 21132 | 82.3 | 25684 |
| Omnitrophia | 73 | 17.3 | 348 | 82.7 | 421 |
| Acetothermiia | 23 | 17.2 | 111 | 82.8 | 134 |
| Cyanobacteriia | 1186 | 16.7 | 5924 | 83.3 | 7110 |
| Parcubacteria | 99 | 14.0 | 607 | 86.0 | 706 |
| **Zetaproteobacteria** | **10** | **8.9** | **102** | **91.1** | **112** |
| **Fimbriimonadia** | **11** | **6.5** | **157** | **93.5** | **168** |
| **JS1** | **17** | **6.2** | **259** | **93.8** | **276** |
| **Gammaproteobacteria** | **1976** | **3.5** | **54340** | **96.5** | **56316** |
| **Magnetococcia** | **4** | **2.2** | **174** | **97.8** | **178** |

#### Movies

**Movie 1**. "Three domains_plotly"; https://doi.org/10.5281/zenodo.17049073
**Movie 2**. "Three domains_polygon_plotly"; https://doi.org/10.5281/zenodo.17049153
**Movie 3**. "Archaea seqspace", "Archaea_polygon_centroid_plotly"; https://doi.org/10.5281/zenodo.17049195
**Movie 4**. "Bacteria seqspace", "Bacteria_polygon_centroid_plotly"; https://doi.org/10.5281/zenodo.17049222
**Movie 5**. "Eukaryota_seqspace", "Eukaryota_polygon_centroid_plotly"; https://doi.org/10.5281/zenodo.17049247

### M10bases Dataset Analysis

#### 1. Dataset creation

In addition to the CAPASYDIS-based analysis of the NR99 dataset which consisted of >330,000 unique, long (>1,200 bases) sequences (see main manuscript), the goal here was to generate a comprehensive collection of DNA sequences that spans the full sequence diversity space of a given sequence of length *n*, meaning that all possible mutations are represented. For n=10 bases, we obtain a set of 1,048,576 distinct sequences ($4^n$). For n=20 bases, this reaches a set containing 1,099,511,627,776, i.e. approximately $1.10\times10^{12}$, distinct sequences, which is far above our computing capacity.
For this demonstration, a go program was created to generate the set of all possible sequence combinations for a 10-base-long sequence with (A,T,G,C) bases. The corresponding dataset and the scripts are available at: https://github.com/RametteLab/CAPASYDIS/tree/main/M10bases

The multiFASTA file "sequences10bases.fasta" obtained after running the script ***01_create_sequences.go*** contains 1,048,576 unique sequences. This is what we call the "M10bases" dataset. The headers of the FASTA file reflect the number of mutations present in each sequence as compared to the initial sequence "AAAAAAAAAA".

The first lines of the "sequences10bases.fasta" file consist of:

```
>S0aaaaa
AAAAAAAAAA
>S1aaaab
TAAAAAAAAA
>S1aaaac
GAAAAAAAAA
>S1aaaad
CAAAAAAAAA
>S1aaaae
ATAAAAAAAA
...
```

where each header of the sequences starts with the letter "S" and is followed by a number, which corresponds to the number of mutations for the given sequence as compared to S0aaaaa. This number is combined with a unique 5-letter identifier to form a unique name for each sequence.
Among the 1,048,576 FASTA sequences, >98% have 5 to 10 mutations:

| | Number of mutations (Nmut) per sequence | | | | | | | | | | |
|---|---|---|---|---|---|---|---|---|---|---|---|
| | 0 | 1 | 2 | 3 | 4 | 5 | 6 | 7 | 8 | 9 | 10 |
| **Sequence number (%)** | 1 (<.01%) | 30 (<.01%) | 405 (.04%) | 3,240 (.31%) | 17,010 (1.62%) | 61,236 (5.84%) | 153,090 (14.60%) | 262,440 (25.03%) | 295,245 (28.16%) | 196,830 (18.77%) | 59,049 (5.63%) |

After applying CAPASYDIS build_axes_0.1.8 script, we identified the sequence "S10atxbh" at position number 349,526 in the MSA as the most distant sequence to S0aaaaa. This is expected given the *delta* table we used (See Table 1 in the manuscript), where A ↔ T was associated with the largest distance in all the pairs involving the adenine base.

```
```{sh}
 build_axes -i inputMSA -R1 1 -max
```
```

Repeating this procedure with S10atxbh to find its most distant sequence, we found sequence S10bnuco.

```
```{sh}
grep -A1 "S10atxbh" inputMSA
```
>S10atxbh
TTTTTTTTTT

```{sh}
build_axes -i inputMSA -R1 349526 -max
```
S10atxbh,349526,S0aaaaa,1,0.6691823746923862

```{sh}
grep -A1 "S0aaaaa" inputMSA
```
```

We obtained a possible system of 3 CAPASYDIS references for our dataset, as follows:

| Name | Sequence position | Base composition |
|---|---|---|
| S0aaaaa | 1 | AAAAAAAAAA |
| S10atxbh | 349,526 | TTTTTTTTTT |
| S10bnuco | 699,051 | GGGGGGGGGG |

Using build_axes, we built CAPASYDIS 2D coordinate axes with (REF1=1, REF2=349526), and then 2D additional axes with (REF1=1, REF2=699051). The csv files of these two outputs were then merged to produce a CAPASYDIS 3D coordinate system (REF1=1, REF2=699051, REF3=349526) using the provided go function *merge3D*.

```
```{sh}
FileR1R2=output_build_axesv0.1.8_R1_As_R2_Ts/output.csv # R1 R2
FileR1R3=output_build_axesv0.1.8_R1_As_R3_Gs/output.csv  # R1 R3
mergedOutput=output_build_axesv0.1.8_R1_R2_R3/output_R1_R2_R3.csv
merge3D -i FileR1R2 -j FileR1R3 -o mergedOutput
wc -l mergedOutput #1048577
head mergedOutput
```
    x,y,z,label
    0,0.6691823747,0.0778113585,S0aaaaa
```

```
    0.1435747169,0.5374491816,0.1690887007,S1aaaab
    0.0182533514,0.6529889579,0.0623373963,S1aaaac
    0.10819395750000001,0.5708622363,0.1989429388,S1aaaad
```

We then used the following "pattern file" (Patterns.tsv) to add colors to the coordinate file:

```
   "S_10" -> saddlebrown
    "S1" ->  red
    "S2" -> deeppink
    "S3" -> magenta
    "S4" -> mediumorchid
    "S5" -> blue
    "S6" -> deepskyblue3
    "S7" -> green3
    "S8" -> palegreen2
    "S9" -> sienna3
```

```
```{sh}
WD=output_build_axesv0.1.8_R1_R2_R3
colorCSVTaxonomy -v # version:0.1.3
inputCSV=WD/output_R1_R2_R3_modif.csv
outputCSV=WD/output_R1_R2_R3_with_color.csv
colorCSVTaxonomy -i inputCSV -o outputCSV -p Patterns.tsv
head outputCSV
```
x,y,z,label,color
0,0.6691823747,0.0778113585,S0aaaaa,black
0.1435747169,0.5374491816,0.1690887007,S1aaaab, red
0.0182533514,0.6529889579,0.0623373963,S1aaaac, red
0.10819395750000001,0.5708622363,0.1989429388,S1aaaad, red
0.10249910620000001,0.5751370243,0.1429788685,S1aaaae, red
0.2460738231,0.4434038313,0.2342562107,S2aaaaf,deeppink
0.1207524576,0.5589436076000001,0.1275049063,S2aaaag,deeppink
0.2106930638,0.476816886,0.2641104488,S2aaaah,deeppink
0.0129231222,0.6575268235,0.0668560057,S1aaaai, red
0.156497839,0.5257936305,0.1581333479,S2aaaaj,deeppink
0.0311764735,0.6413334068000001,0.0513820435,S2aaaak,deeppink
0.1211170797,0.5592066852,0.187987586,S2aaaal,deeppink
0.0770618426,0.5988221047,0.1644909682,S1aaaam, red
0.2206365595,0.4670889116,0.25576831040000003,S2aaaan,deeppink
0.095315194,0.5826286879,0.149017006,S2aaaao,deeppink
0.18525580020000001,0.5005019663,0.2856225485,S2aaaap,deeppink
0.0839627965,0.5921445262,0.1311943571,S1aaaaq, red
0.2275375134,0.4604113332,0.22247169930000002,S2aaaar,deeppink
0.1022161479,0.5759511094,0.1157203948,S2aaaas,deeppink
…
```
```

## 2. CAPASYDIS analyses

The detailed analyses of the 2D and 3D CAPASYDIS results are presented below. The scripts are available at: https://github.com/RametteLab/CAPASYDIS/tree/main/M10bases/03_R_analyses

### How numerically distinct are the CAPASYDIS coordinates?

If we consider each single CAPASYDIS axis on its own, we found the x axis (with 1,048,508 numerically distinct values), the y axis (1,048,476 distinct values), and z axis (1,048,469 distinct values) to consist of >99.99% unique 1D coordinate values, among the 1,048,576 sequences of the full dataset. After creating the 2D (x,y) or 3D (x,y,z) coordinates, then all sequence
```

coordinates became 100% numerically distinct from each other. This observation was valid when using either the forward or reverse reading directions of the sequences in the MSA. Given the way the axes were calculated (i.e. as the maximum distance to the first reference, and repeating this calculation for the second axis), the basic statistics for the axis coordinates are identical for both read directions (forward and reverse), as S0aaaaa is symmetric and consist of the base "A" repeated 10 times. Here only the forward CAPASYDIS coordinate data is presented (truncated to 4 decimals):

| | X axis | Y axis | Z axis |
|---|---|---|---|
| **N values** | 1,048,576 | 1,048,576 | 1,048,576 |
| **Minimum** | 0.0000 | 0.0000 | 0.0000 |
| **1st quantile** | 0.2692 | 0.2895 | 0.2577 |
| **Median** | 0.3416 | 0.3566 | 0.3280 |
| **Mean** | 0.3423 | 0.3559 | 0.3286 |
| **3rd quantile** | 0.4147 | 0.4229 | 0.3990 |
| **Maximum** | 0.7293 | 0.6692 | 0.6949 |

The scatterplots below illustrate the 2D visualization of the M10bases dataset for the 100, 1,000, 10,000 first sequences combined with their associated number of mutations (Nmut):

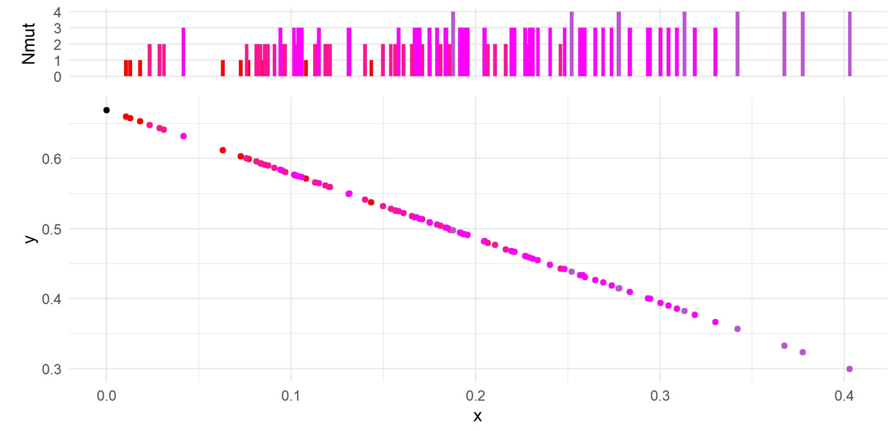

**Scatterplots of the CAPASYDIS 2D (x,y) coordinates of the M10bases dataset for the <u>first 100 sequences (forward direction)</u>.** The marginal plot displays the number of associated mutations (Nmut) for each sequence (each dot) of the x axis.

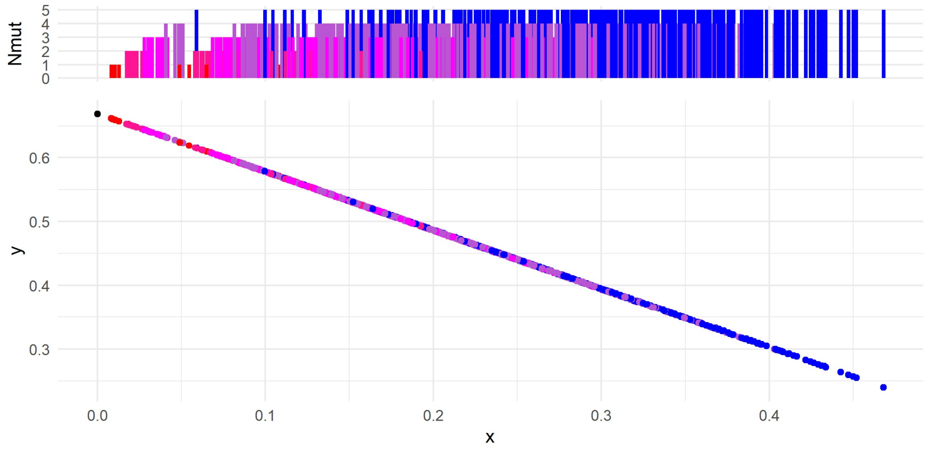

**Scatterplots of the CAPASYDIS 2D (x,y) coordinates for the M10bases dataset for the <u>first 1,000 sequences</u>.** The marginal plot displays the number of associated mutations (Nmut) for each sequence (each dot) of the x axis.

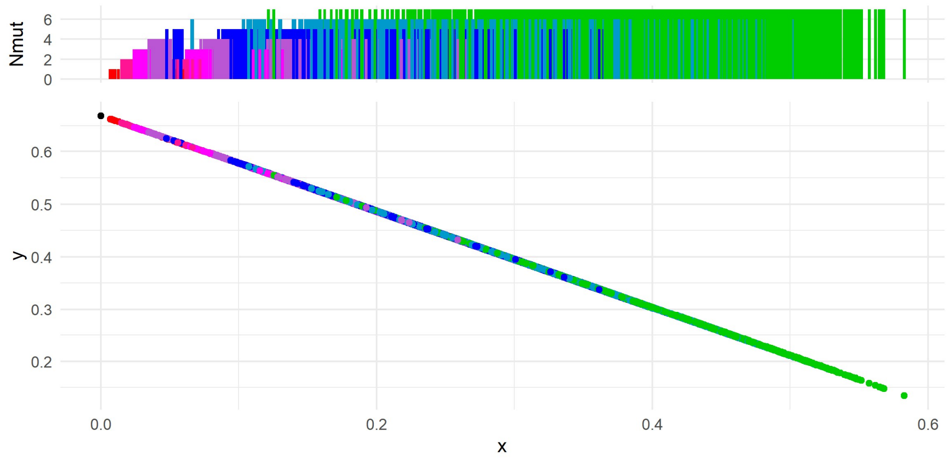

**Scatterplots of the CAPASYDIS 2D (x,y) coordinates for the M10bases dataset for the <u>first 10,000 sequences</u>.** The marginal plot displays the number of associated mutations (Nmut) for each sequence (each dot) of the x axis.

**Conclusion**: The 2D representation of the M10bases dataset is consistent with a linear gradient of the mutation landscape from low to high mutated sequences along the seqverse.

#### CAPASYDIS 3D visualization

<table>
<tr><td>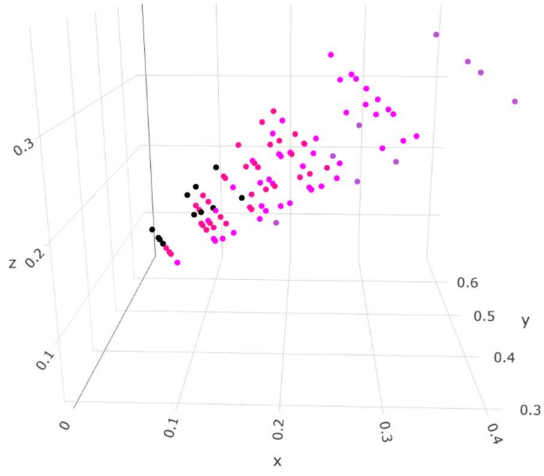
</td><td>The top 100 sequences are displayed.</td></tr>
<tr><td>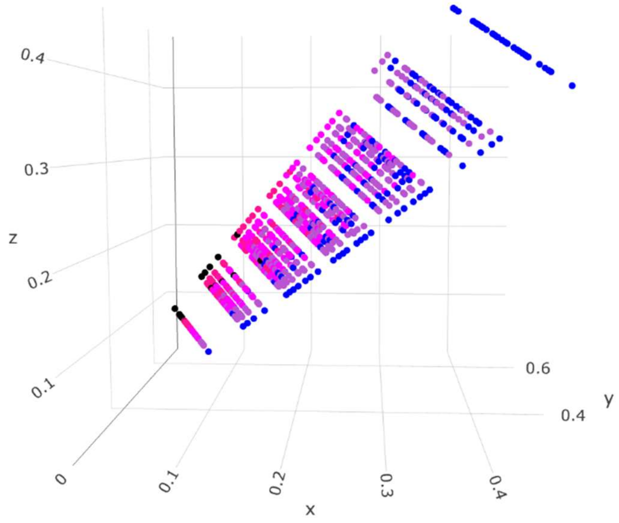
</td><td>The top 1,000 sequences are displayed.</td></tr>
<tr><td>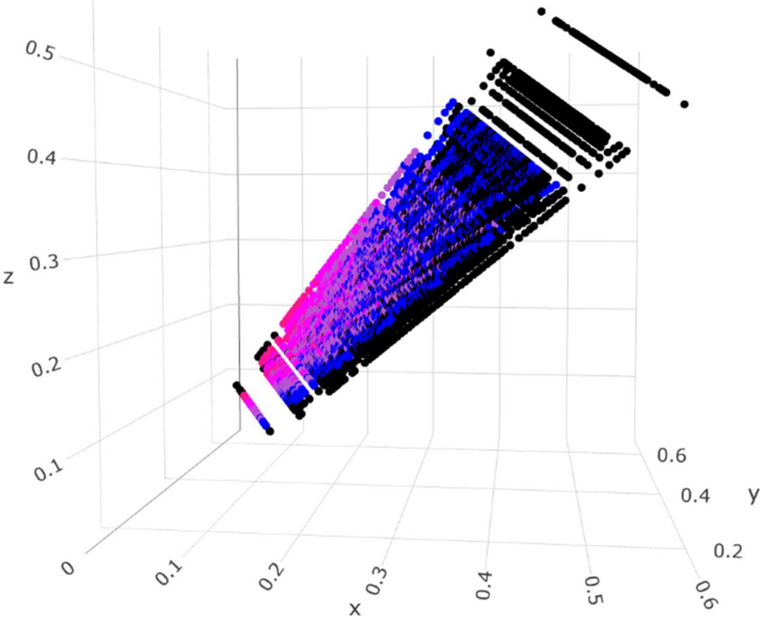
</td><td>The top 10,000 sequences are displayed.</td></tr>
</table>

| | |
|---|---|
| 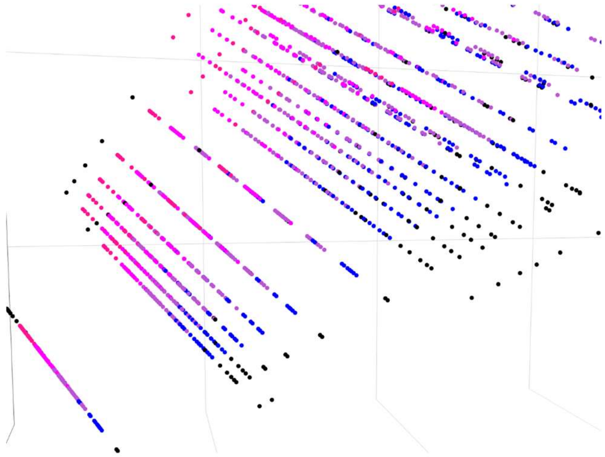 | Closer look at the top 10,000 sequences, illustrating the uniqueness of each point. |
| 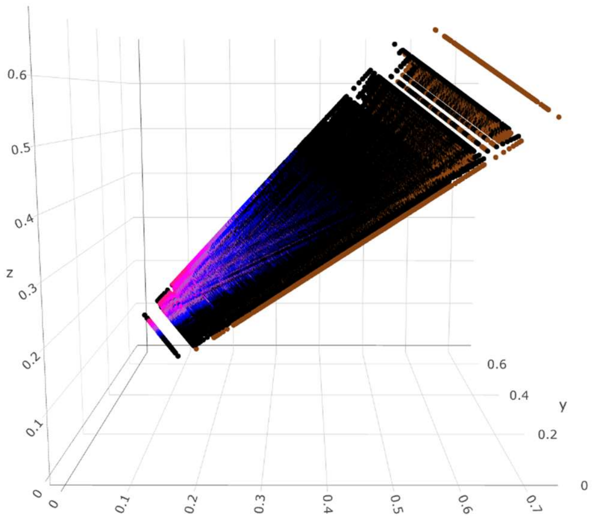 | A visualization of the top **500,000** sequences. Beyond this point, the interactive rendering using plotly() in R did not manage to cope with the very large number of points using a standard laptop device. Switching to the provided D3.js scripts to display larger datasets is recommended (https://github.com/RametteLab/CAPASYDIS/tree/main/JavaScript). |
| **A** **B** 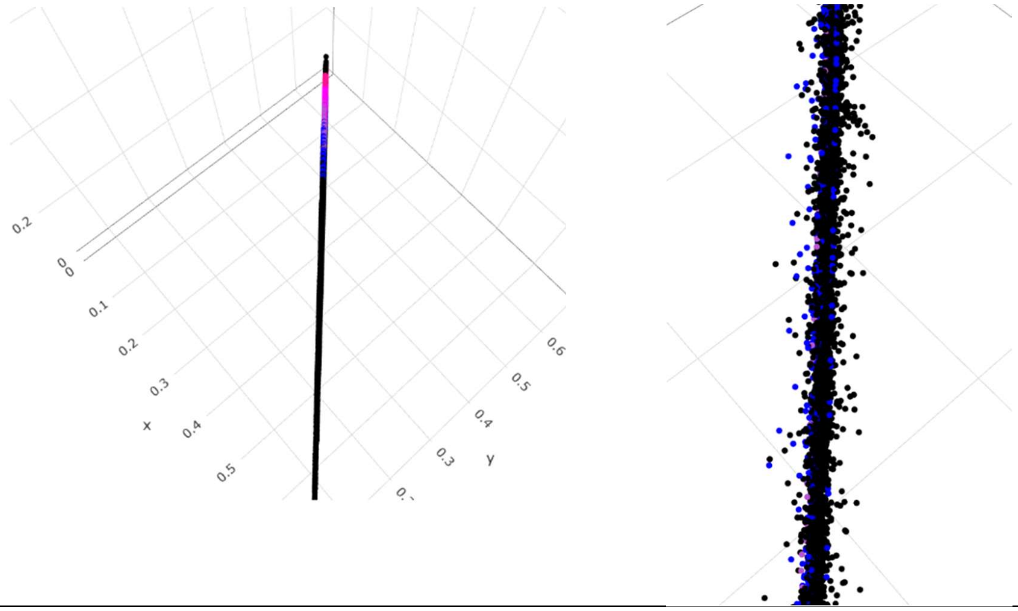 | |
| **A**) Visualization of the top **500,000** sequences in 3D. When viewed from the z axis perspective, the seqverse seems to consist of a flat surface. **B**) A closer look at the edge of this "flat" surface shows that sequences are close, yet distinctly positioned around this "plane". | |

**Conclusion**: A particular diversity plane was evidenced when exploring the full diversity landscape of the 10-base-long sequence seqverse. The 3D visualization of the M10bases dataset further supports the idea that each sequence is distinctly represented using CAPASYDIS at a definitive and orderly position. For this dataset, as for the NR99 dataset, the smallest difference between any two adjacent sequence scores is still much larger than our chosen numerical precision ($10^{-10}$), and millions of times larger than the absolute threshold for a computational collision (machine epsilon $2.2\times10^{-16}$). Consequently, the sequences represented in the seqverses are perfectly resolvable.

#### Comparison of CAPASYDIS coordinates for 5' vs. 3' sequence directions

The sequences in "sequences10bases.fasta" were reversed (e.g. the sequence "ACTGACTGAC" was changed into "CAGTCAGTCA" in the reversed dataset). Then the same CAPASYDIS coordinate calculations were undertaken using CAPASYDIS build_axes_0.1.8 script, as described previously (section "1. Dataset creation").

The coordinates of the forward and reverse datasets were merged, and below is a snapshot at the top 6 entries of that merged table, with "F" standing for the data originating from the sequences in the forward direction, and "R_" for those in the reverse reading direction:

| | F_x | R_x | F_y | R_y | F_z | R_z | F_label | R_label | F_color | R_color | F_Nmut | R_Nmut |
|---|---|---|---|---|---|---|---|---|---|---|---|---|
| 1 | 0.00000000 | 0.000000000 | 0.6691824 | 0.6691824 | 0.07781136 | 0.07781136 | S0aaaaa | S0aaaaa_R | black | black | 0 | 0 |
| 2 | 0.14357472 | 0.046200672 | 0.5374492 | 0.6267922 | 0.16908870 | 0.10718570 | S1aaaab | S1aaaab_R | red | red | 1 | 1 |
| 3 | 0.01825335 | 0.005785165 | 0.6529890 | 0.6638928 | 0.06233740 | 0.07290708 | S1aaaac | S1aaaac_R | red | red | 1 | 1 |
| 4 | 0.10819396 | 0.034667743 | 0.5708622 | 0.6374056 | 0.19894294 | 0.11695769 | S1aaaad | S1aaaad_R | red | red | 1 | 1 |
| 5 | 0.10249911 | 0.048689031 | 0.5751370 | 0.6245090 | 0.14297887 | 0.10876779 | S1aaaae | S1aaaae_R | red | red | 1 | 1 |
| 6 | 0.24607382 | 0.094889704 | 0.4434038 | 0.5821188 | 0.23425621 | 0.13814213 | S2aaaaf | S2aaaaf_R | deeppink | deeppink | 2 | 2 |

...

The visual comparisons of the CAPASYDIS 2D (x,y) coordinates for the M10bases dataset for both the forward and reverse sequence directions indicate very similar seqverse shapes:

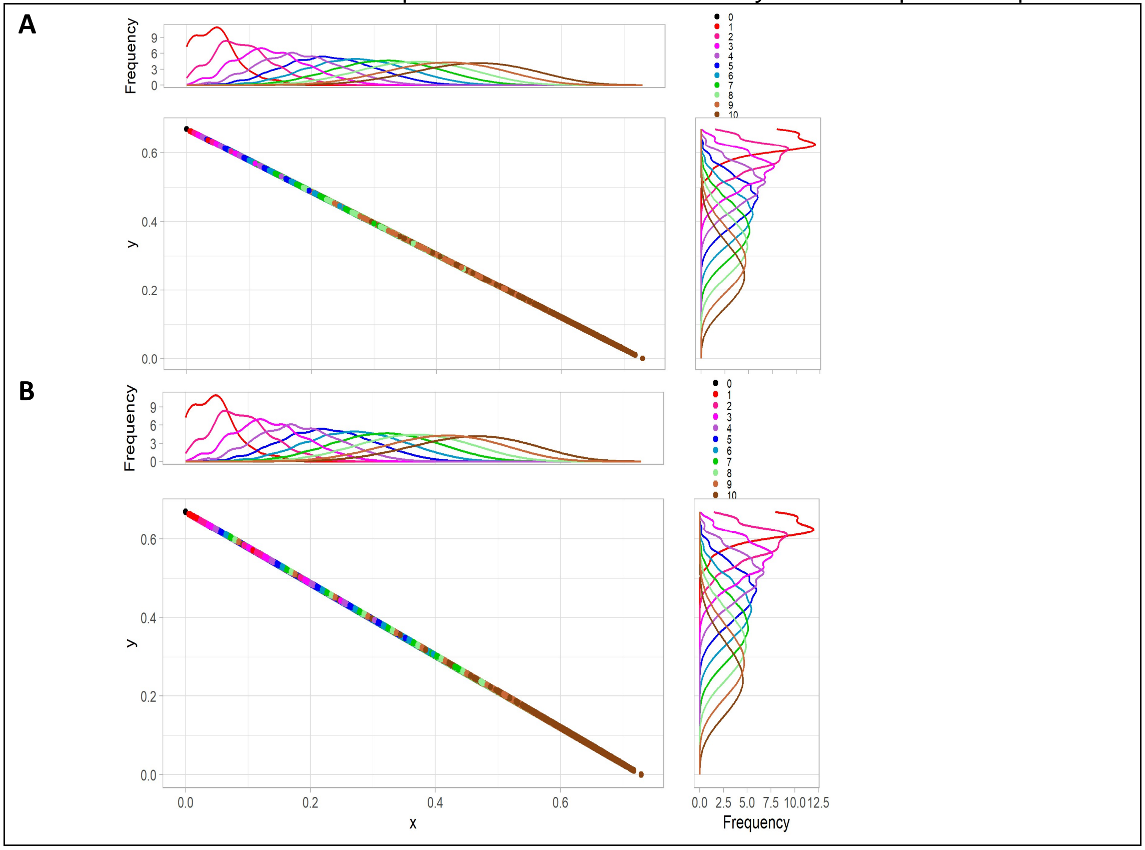

**Scatterplots of the CAPASYDIS 2D (x,y) coordinates for the M10bases dataset for both the forward and reverse sequence directions.** The marginal plots display the frequency distribution of the number of associated mutations (colors indicate Nmut) for each sequence (one dot is one sequence). **A**) Forward direction of the sequences using coordinates (F_x, F_y). **B**) Reverse reading direction of the same set of sequences using coordinates (R_x, R_y).

Beyond the (indirect) visual comparisons of the 2D seqverses shown above, the next step aimed at quantifying the degree of correspondence between the seqverses based on the forward vs. reverse sequence directions. If the coordinate system is symmetric, we would expect a strong correlation between the seqverses, such that a simple rotation of one seqverse coordinate system would align it with the other. We thus calculated the linear correlation coefficients between the seqverses in 1D, 2D and 3D of the forward and reverse datasets, and obtained:

| Seqverse structure | Coordinate comparison (X vs. Y) | Correlation coefficient |
|---|---|---|
| **1D** | [F_x] vs. [R_x] | Pearson's r = 0.776 (P<0.001) |
| **1D** | [F_y] vs. [R_y] | Pearson's r = 0.776 (P<0.001) |
| **1D** | [F_z] vs. [R_z] | Pearson's r = 0.776 (P<0.001) |
| **2D** | [F_x, F_y] vs. [R_x, R_y] | RV = 0.6024 |
| **3D** | [F_x, F_y, F_z] vs. [R_x, R_y, R_z] | RV = 0.6025 |

For the 2D and 3D coordinate tables, we determined how similar the *multivariate structures* of forward and reverse datasets were by using the RV coefficient (Robert and Escoufier 1976), implemented in FactoMineR (https://cran.r-project.org/web/packages/FactoMineR/index.html). This function takes two multivariate tables (X and Y) with the same number of rows (here the sequences) and computes the RV coefficient, a multivariate analogue of a linear correlation between data tables, based on the covariance (or cross-product) matrices.

As a complement to the correlation table provided above, we illustrated the pairwise visual comparisons of the sequence coordinates originating from the forward vs. reverse datasets for each axis below. Visual comparisons of the 2D and 3D objects were not conducted due to their high computational cost.

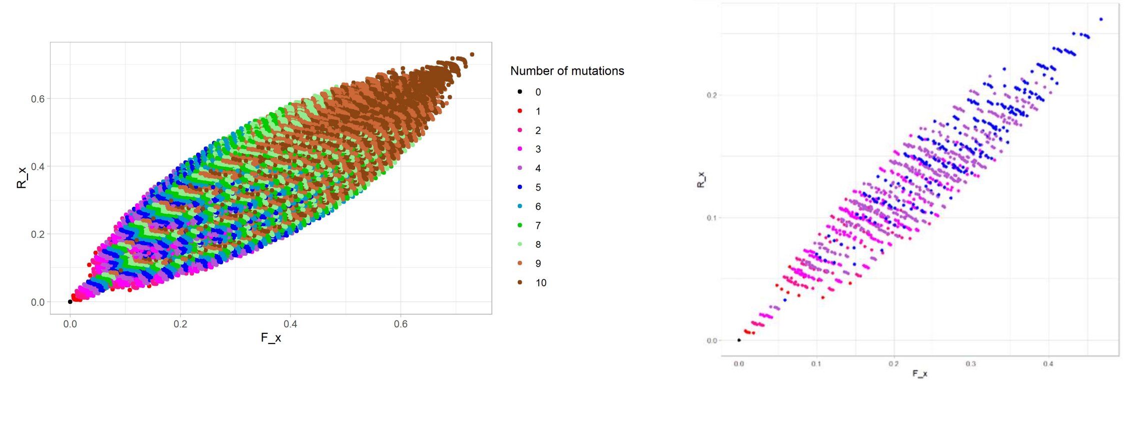

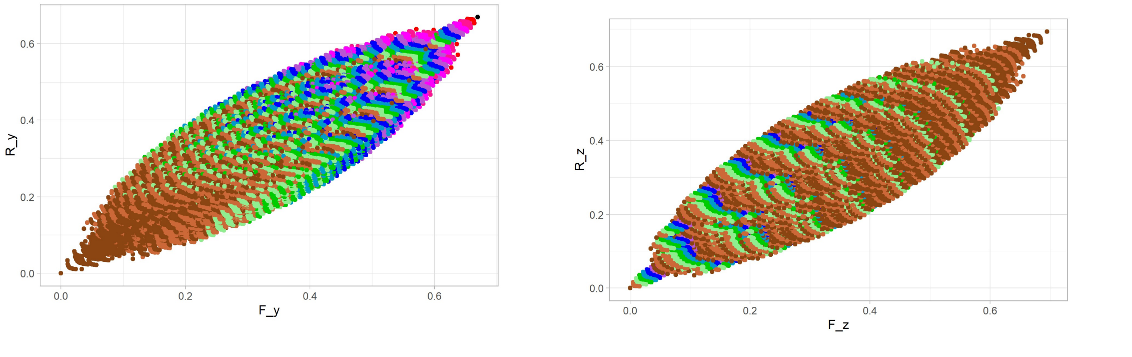

**Scatterplots of each CAPASYDIS 1D coordinate comparing the forward ("F_") and reverse ("R_") coordinates for the whole M10bases dataset.** The color of each sequence follows the same scheme as in the previous figures. The top right panel is a plot of the top 1,000 sequences of the (F_x, R_x) plot.

**Conclusion**: The seqverse representations using either the forward or reverse sequence directions of the same dataset are simply not "mirror" images of each other, and cannot be derived by simply rotating one seqverse coordinate to produce the other representation. Although the forward-based and reverse-based sequence seqverses consist of distinct dataset structures, they are correlated to each other, with linear correlation coefficients ranging from 0.60 (2D, 3D) to 0.78 (1D). This behavior is expected given the asymmetric nature of the CAPASYDIS distance calculations, which incorporate positional information from the sequences.

#### 3. Comparison of CAPASYDIS to alternative approaches

Similarly to the detailed comparative analyses of the *E. coli* seqverse vs. standard phylogenetic approaches (Figure 7 in the manuscript), alternative analytical approaches to CAPASYDIS are tested here on the M10bases dataset.

For this example, we selected the top 100 sequences of M10bases to create a dataset that is easily visualizable via alternative approaches. Phylogenetic tree computations were done with iqtree (see main manuscript and GitHub repository for additional information), using the following parameters:

````
```{sh}
iqtree -s  MSAtop100.fasta -m TIM2+F+I -nt 60
```
````

Rate parameters:  A-C: 0.00124  A-G: 0.00010  A-T: 0.00124  C-G: 1.00000  C-T: 0.39326  G-T: 1.00000
Base frequencies:  A: 0.749  C: 0.065  G: 0.069  T: 0.117
Proportion of invariable sites: 0.565

Further, the ML distances to the reference sequence (S0aaaaa) consist of only 6 distinct numerical values, which are summarized in the following table (the maximum numerical precision was used here):

| Classical phylogenetic distances to the reference | Number of sequences matching this distance value |
|---|---|
| 0.0000000 | 1 |
| 8.7451681 | 7 |
| 8.9999987 | 17 |
| 8.9999982 | 18 |
| 8.9999989 | 3 |
| 9.0000000 | 54 |

In comparison, we compared the top 100 asymdist values from the CAPASYDIS file "output_R1_R2_R3_with_color.csv" to these classical phylogenetic distances (iqtree.mldist from the iqtree output). All 100 sequences produced 100 distinct coordinates via CAPASYDIS (1D, 2D or 3D), as illustrated below (only 1D seqverse is shown), further highlighting the advantage of using CAPASYDIS, which produces distinct and definitive coordinates for each distinct sequence. In addition, the approach is computationally scalable, which allows a standardized exploration of the seqverses of datasets of various complexity and sizes:

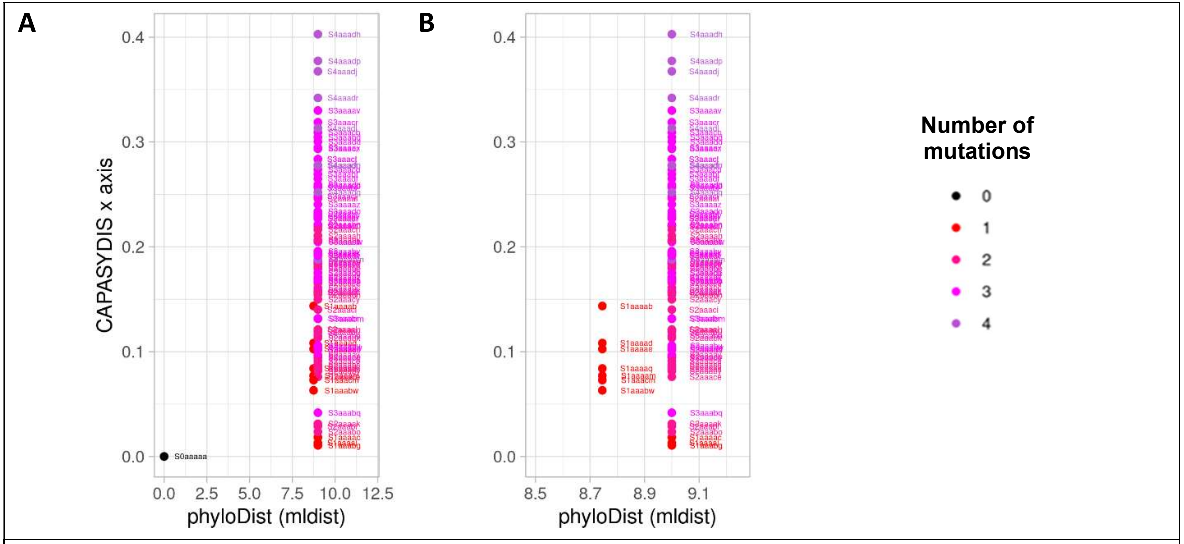

**A**) CAPASYDIS coordinates (x axis only) plotted vs. classical phylogenetic distances (MLDIST). The color code of each dot (one sequence) reflects the number of mutations in each sequence. **B**) Closer view for MLDIST values 8.5 to 9.5.

As visualized using classical phylograms below, the 100 sequences represent a diverse set of sequences, as expected given the way they were designed.

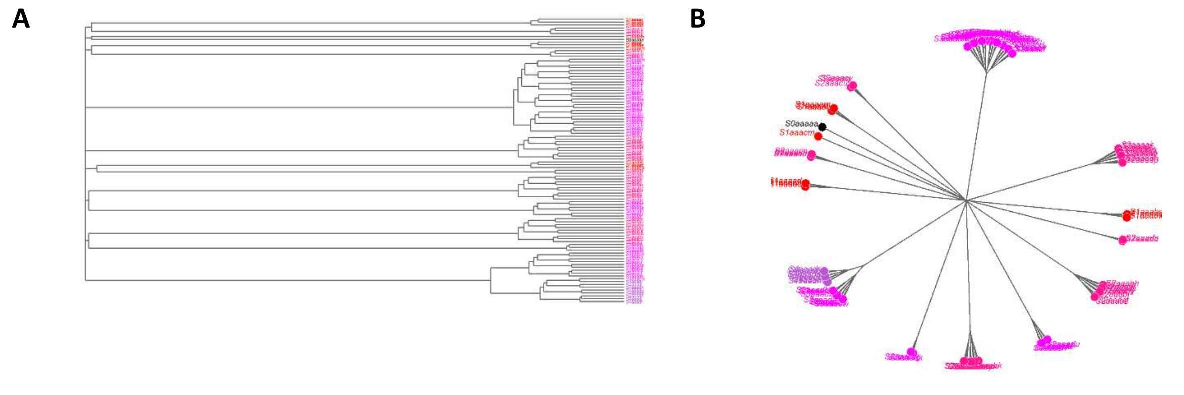

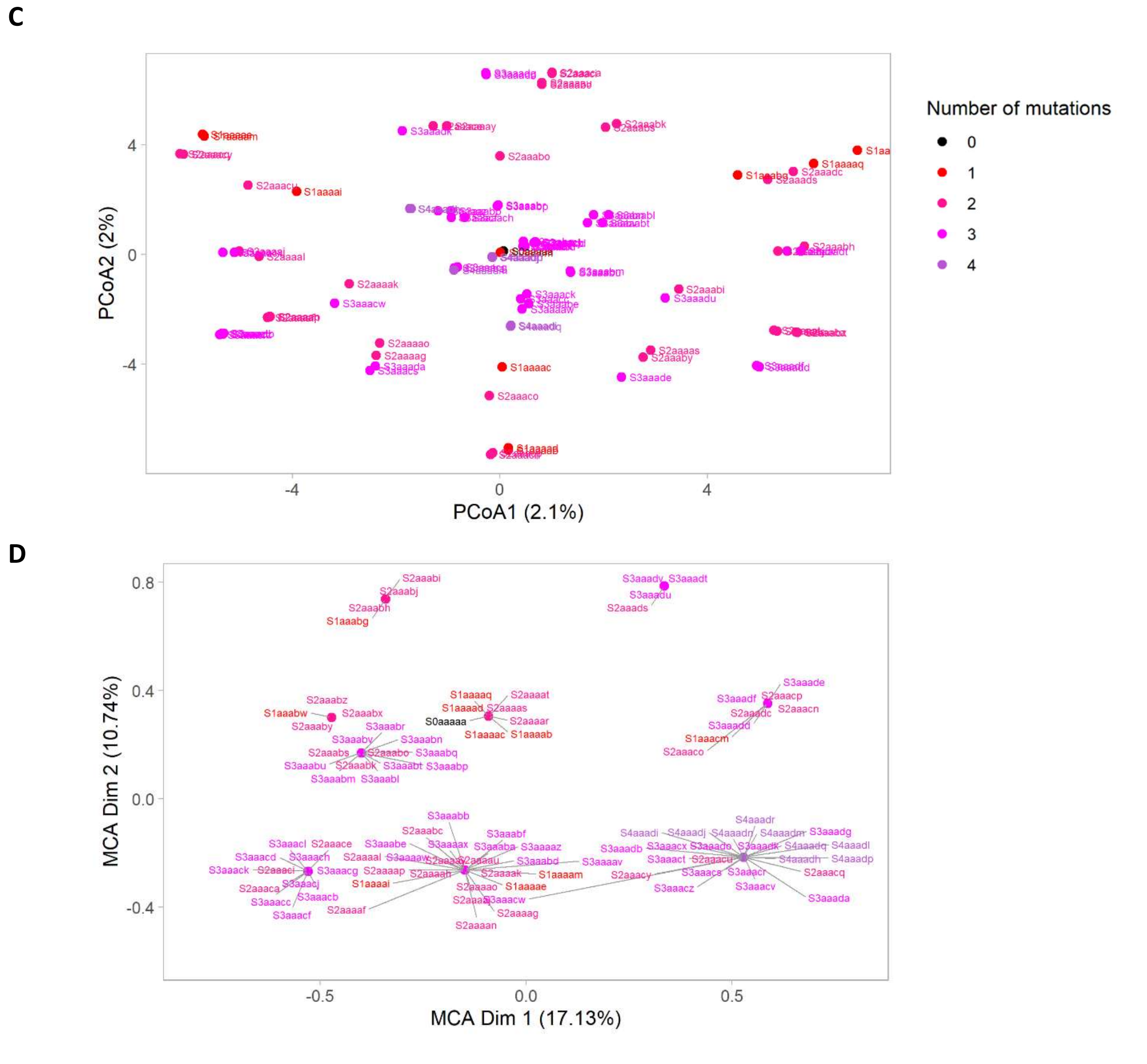

**Alternative methods to represent the diversity in M10bases. A**) Classical phylogram based on the maximum likelihood (ML) distances. **B**) Unrooted representation of the phylogram shown in A). **C**) Visualization of the ML distances using Principal Coordinate Analysis (PCoA) using the function *wcmdscale* implemented in the R library vegan. **D**) Multiple Correspondence Analysis (MCA) ordination derived from the singular value decomposition (SVD) of a specially transformed, center-scaled indicator matrix generated via constant-length one-hot encoding of the M10bases MSA. The color coding of the labels in all figures is consistent with the provided legend (Nmut).

Because standard **phylograms** compress multidimensional sequence data into a one-dimensional branching layout, classical Principal Coordinate Analysis (PCoA) was subsequently used to project the pairwise ML distance matrix into a continuous spatial coordinate system. The classical **PCoA** dimensionality reduction approach proved highly ineffective at representing the true high-dimensional nature of the small but complex sequence space, as the first three ordination axes explained only a remarkably small percentage of the total variance (5.6% in total). In a successful ordination, the first two axes typically capture a substantial chunk of the total variation—often ranging from 40% to over 70%—which allows researchers to look at a 2D plot and confidently trust that visual distance accurately reflects true dissimilarity. Because the underlying variance is here scattered across hundreds of minor axes rather than concentrated into a few major gradients, a standard low-dimensional metric projection cannot reliably summarize the sample relationships in a visual space.

Finally, we applied Multiple Correspondence Analysis (MCA), an ordination technique derived from the singular value decomposition (SVD) of transformed categorical data. MCA and PCoA differ fundamentally in their input data requirements: PCoA requires a pre-calculated pairwise distance matrix (here we used the ML distance matrix), which is subsequently projected into a continuous coordinate system via Eigenvalue Decomposition. Conversely, MCA completely bypasses traditional distance calculations, applying a Chi-squared transformation directly to a rectangular 0/1 indicator matrix generated by one-hot encoding. This mathematically weights features by their rarity so that unique sequence mutations carry more statistical influence than common ones—before using SVD to map the variances of both shared presences and absences simultaneously (Greenacre 2007).

When applied to the M10bases dataset, a major limitation of MCA was yet revealed: Uninformative columns containing either all zeros or all ones must be meticulously removed prior to analysis, as these zero-variance variables cause a division-by-zero error during the Chi-squared standardization step and will crash the algorithm. Therefore, the initial MSA contained 10 base positions, and after one-hot encoding (see GitHub repository, file "05_Analyses_PCoA_MCA.R", we obtained 40 binary columns encoding the presence or absence of each base {A,T,G,C}. After removing constant columns, we ended up with only 14 valid columns. This architectural dependency creates a significant scalability bottleneck: Because the feature space must be dynamically adjusted to strip out zero-variance columns each time new mutation data is introduced, the underlying dimensions of the MCA model change, preventing the direct projection of new sequences onto a fixed, historical reference coordinate system.

The first three MCA dimensions captured a cumulative total of 38.38% of the dataset's variance, with Dimension 1, 2, and 3 accounting for 17.13%, 10.74%, and 10.50% of the total inertia, respectively. Importantly, if we use a numerical precision of e-10, only 9 unique values were obtained on each MCA axis. If we allowed the maximum machine precision we only obtained 21 and 51 distinct values for MCA axis 1 and 2 respectively, for the 100 different sequences.

**Overall conclusion for the comparative analysis of the M10bases dataset**
Using the exhaustive M10bases dataset of all 1,048,576 possible 10-bp sequences, CAPASYDIS assigned each sequence to unique, numerically stable 2D/3D coordinates that form orderly mutation gradients and a well-defined diversity plane, achieving full resolvability far beyond numerical precision limits. In stark contrast, classical maximum likelihood (ML) distances collapse the top 100 sequences onto just 6 highly compressed, discrete numerical values, demonstrating that standard phylogenetic solutions are not unique and fail to well capture fine sequence details and specific mutation patterns. Application of traditional ordination techniques revealed severe architectural constraints: Classical PCoA proved highly ineffective, scattering variance across hundreds of minor gradients and capturing a remarkably low 5.6% of the total variance across its first three axes. While MCA captured a higher cumulative variance of 38.38% on its first three dimensions, it introduced significant computational limitations: it failed to provide individual resolution for the sequences—yielding as few as 9 unique values per axis at standard precision—and required the meticulous stripping of zero-variance columns (reducing the one-hot encoded matrix from 40 to just 14 valid features). This

structural dependency creates a critical scalability bottleneck, as the MCA feature space must be dynamically redefined each time new mutation data enters the analysis, preventing sequence projection onto a fixed historical reference. Furthermore, while forward and reverse CAPASYDIS representations are strongly but imperfectly correlated—confirming they are not simple mirror images—the algorithm natively incorporates positional sequence context. Ultimately, while traditional frameworks struggle with coordinate duplication or dynamic feature constraints, CAPASYDIS offers a continuous, scalable coordinate space that preserves unique sequence resolution across large datasets.

#### Cited literature

Greenacre, M., 2007. *Correspondence Analysis in Practice*. 2nd ed. Boca Raton: Chapman and Hall/CRC. https://doi.org/10.1201/9781420011234

Robert, P., and Y. Escoufier. "A Unifying Tool for Linear Multivariate Statistical Methods: The RV-Coefficient." *Journal of the Royal Statistical Society. Series C (Applied Statistics)*, vol. 25, no. 3, 1976, pp. 257–65. *JSTOR*, https://doi.org/10.2307/2347233

#### Algebraic Independence and Exhaustive Matrix Audit of Score Uniqueness in 1D and 2D CAPASYDIS representations

Coordinate uniqueness is at the core of the CAPASYDIS approach, as the main idea is that each query sequence of a MSA is associated to a unique and fixed Euclidean coordinate in 1D, 2D or 3D seqverses. As sequence length n increases, the density of the projection space (the "seqverse") also increases, as illustrated in Figure 2 of the main manuscript. Even if we did not encounter numerical collisions in our 2D and 3D seqverses when using the large biological NR99 dataset and the complex M10bases dataset, the main question is whether specific combinations of mutation patterns could "almost coincide" and produce identical scores even if the sequences have little similarities. This fundamental concern regarding structural uniqueness and numerical resolution is rigorously addressed through the following two points:

**1. Mathematical uniqueness.** The concern regarding "colliding" points is addressed by fundamental number theory. For two different mutation patterns to produce identical scores, their respective sums of radicals would need to be **rationally dependent**. Two terms are rationally dependent if one can be transformed into another by a rational multiplier (e.g., $\sqrt{18} = 3\sqrt{2}$). In the CAPASYDIS framework, sequence positions $i$ are integers and delta values take only a limited number of constant values. The resulting radicands (e.g., 1+delta, 2+delta, 3+delta,...) are **square-free relative to each other**, meaning they do not share a common radical "base". This independence is guaranteed by the **Besicovitch Theorem (1940)**, which proves that such radicals are linearly independent over the field of rational numbers $\mathbb{Q}$. Consequently, each mutation pattern produces a mathematically unique irrational sum that functions as a numerical "fingerprint". No combination of mutations at different positions can perfectly "mimic" the sum of another set of positions.

Because CAPASYDIS allows position-specific shifts ($\text{delta}_i$) to vary dynamically along a sequence, as proposed in Table 1 (main manuscript), validating a single static sequence configuration is insufficient to prove the framework's overall stability. A structural vulnerability would emerge if a mutation mapping at Position A utilizing $\text{delta}_m$ could yield the exact same underlying square-free core as Position B utilizing $\text{delta}_n$. If such a shared algebraic base existed, distinct multi-position mutational patterns could linearly cancel each other out or perfectly mimic one another, resulting in an analytical "collision" where different biological sequences produce identical mathematical scores.

Performing an exhaustive matrix audit of all possible position-delta building blocks is therefore essential to rule out these structural dependencies. According to fundamental field theory established by Besicovitch (1940), a set of real radicals is strictly linearly independent over the field of rational numbers $\mathbb{Q}$ if their integer radicands share no common square-free factors. By executing a global matrix audit across the entire 10 times 10 space of positions (i in {1, ..., 10}) and dynamic offsets ($\text{delta}_j$ in {0.01, 0.02, ..., 0.14}), we verify whether all 100 available building blocks reduce to unique, distinct square-free integer cores.

To prove mathematical uniqueness under this dynamic setup, we must ensure that **no two positions can ever generate the same square-free core, regardless of which delta value they select.** Because a mutation pattern is a summation of these positional roots, a structural collision can only happen if a radicand at Position A using $\text{delta}_m$ shares a square-free core with

Position B using $delta_n$. To test this, we calculate a 10 times 10 matrix of all possible integer radicands:

$$R_{i,j} = 100 . (i + delta_j),$$

where i is the position (1 to 10) and $delta_j$ is the chosen delta value from our set (Table 1, main manuscript). We then extract the square-free core for all 100 possible combinations. If all 100 cores are completely unique, the system is mathematically bulletproof across all possible dynamic assignments.

The script (available at https://github.com/RametteLab/CAPASYDIS/blob/main/Golang/audit_collision/audit_collision_1D.go) builds the entire 10 x 10 interaction matrix, extracts the square-free cores, and checks for **cross-position collisions**. The analysis of all possible position-delta combinations indicated that 7 potential structural dependencies could be encountered when performing the analysis in 1D:

| Combination A | Combination B | Square-Free Core |
|---|---|---|
| Pos 1 with delta=0.02 | Pos 4 with delta=0.08 | 102 |
| Pos 1 with delta=0.08 | Pos 5 with delta=0.07 | 3 |
| Pos 3 with delta=0.12 | Pos 7 with delta=0.02 | 78 |
| Pos 2 with delta=0.02 | Pos 8 with delta=0.08 | 202 |
| Pos 1 with delta=0.01 | Pos 9 with delta=0.09 | 101 |
| Pos 5 with delta=0.13 | Pos 9 with delta=0.12 | 57 |
| Pos 1 with delta=0.12 | Pos 10 with delta=0.08 | 7 |

For instance, let's take the first line of the above table and break it down into its exact arithmetic components. The first line means that in a single-dimension (1D) projection, a mutation at Position 1 and a mutation at Position 4 can share the exact same mathematical "base root" of 102. Because they share a base root, they are rationally dependent, which violates the strict conditions required for 1D uniqueness under Besicovitch's Theorem. This is how this is obtained:

*Combination A (Pos 1, delta = 0.02):*

Score for position 1 = $\sqrt{1 + 0.02}$ = $\sqrt{1.02}$ = $\sqrt{\frac{102}{100}} = \frac{1}{10}\sqrt{102}$

*Combination B (Pos 4, delta = 0.08):*

Score for position 4 = $\sqrt{4 + 0.08}$ = $\sqrt{4.08}$ = $\sqrt{\frac{408}{100}} = \frac{\sqrt{408}}{10} = \frac{\sqrt{4\times 102}}{10} = \frac{2}{10}\sqrt{102}$

We simplified the integer numerators (102 and 408) by pulling out any perfect square factors to find their underlying "square-free core". For **102**, the prime factors are 2×3×17. There are no repeating factors (no perfect squares), so it cannot be simplified any further. Its square-free core is 102. When we look at the final simplified fractions, the structural overlap becomes obvious, because they share the exact same irrational base ($\sqrt{102}$), **position 4 is exactly double the value of position 1**. Because one is a perfect rational multiple of the other, they are linearly dependent over Q. This indicates that a 1D sequence projection space with the specific delta table values may be vulnerable to geometric overlapping, meaning identical coordinates may be obtained for sequences harboring separate mutations at different positions.

However, **the multi-dimensional approach (2D or 3D axes) changes everything.** In a multi-dimensional CAPASYDIS projection, each sequence is mapped to a spatial coordinate vector (e.g., (X, Y) in 2D or (X, Y, Z) in 3D), where each axis calculates its distance relative to a **different reference point**. For a collision to occur in higher dimensions, two different position-delta combinations must collide **on all axes simultaneously**. If Combination A and Combination B collide on Axis X, but map to completely different values on Axis Y, their physical positions in the multi-dimensional space remain uniquely separated.

We thus created a go script **for Higher-Dimensional Audit ("audit_collision_2D.go")** to evaluate a **2D projection system**, where each position-delta combination is evaluated using two separate reference metrics ($R_X$ and $R_Y$). We processed as follows:

1. Compute the square-free cores for Axis X.
2. Compute the square-free cores for Axis Y.
3. Merge them into a coordinate core pair ($Core_X$, $Core_Y$).
4. Check if this exact **ordered pair** repeats anywhere else in the matrix.

Axis X represents our standard base configuration. It calculates the radicand directly from the natural left-to-right position of the sequence:

$Radicand_X$ = position + delta

The second axis simulates a secondary reference point by inverting the positional distance calculation:

$Radicand_Y$ = (11 - position) + delta

For Position 1, Axis Y treats it as being a distance of 10 units away from the opposite end (11 - 1 = 10). For Position 4, it treats it as being 7 units away (11 - 4 = 7).

By pairing these two calculations, every position-delta combination receives a dual geometric identity: a coordinate pair of square-free cores ($Core_X$, $Core_Y$).

For example, let's look at how the script completely breaks the 1D collision between **Pos 1 (delta=0.02)** and **Pos 4 (delta=0.08)** again:

| Combination | Axis X Core | Axis Y Core | Final 2D Coordinate Signature |
|---|---|---|---|
| Pos 1 with delta=0.02 | 102 | 1002 | (102, 1002) |
| Pos 4 with delta=0.08 | 102 | 177 | (102, 177) |

For Pos 1, the raw radicand becomes 10 + 0.02 = 10.02. As we calculated in the previous step, clearing the decimals yields $\sqrt{\frac{1002}{100}}$, i.e. a square-free core of 1002. For Pos 4, the raw radicand becomes 7 + 0.08 = 7.08. Clearing the decimals yields $\sqrt{\frac{708}{100}}$, i.e. a square-free core of 177.

Thus, even though the two positions share a core on Axis X (102), their Axis Y cores are entirely distinct (1002 vs 177). Because the combined pairs (102, 1002) and (102, 177) do not match, **they do not collide in 2D space.**

**After running the go script for the 2D collision audit test, we found zero multi-dimensional coordinate collisions**. Consequently, the "failure" of the audit test for collisions in 1D actually **perfectly proves why a multi-dimensional approach is necessary in CAPASYDIS**.

Note that with a biological dataset, the secondary axis doesn't just invert the position index numbers; it alters the *metric baseline* based on actual sequence characters. If the second axis uses a completely different biological reference sequence, as it is the case in CAPASYDIS suggested usage, the 2D audit script is a conservative proxy—meaning that real-world biological variation will break these 1D ties even more aggressively than the simple mathematical inversion used in the script.

The algebraic results obtained with the 1D and 2D audit scripts are also supported by the empirical results obtained when applying CAPASYDIS to the fully mutated M10bases dataset, where few cases (<0.01%) of non-unique coordinates for the 1D representation of the full dataset were identified, for the x axis (with 1,048,508/1,048,576 numerically distinct values), the y axis (1,048,476/1,048,576 distinct values), and z axis (1,048,469/1,048,576 distinct values). After creating the 2D (x,y) or 3D (x,y,z) coordinates, then all sequence coordinates became 100% numerically unique.

While a brute-force search across the complete sequence M10bases dataset, comprising 4^10 = 1,048,576 individual sequences, empirically confirms that no numerical collisions occurred within that specific dataset, our global audit scripts provide a definitive mathematical proof that covers the entire space without needing to generate it. The matrix audit analyzes the fundamental 100 position-delta building blocks that are used to construct those millions of final scores in any dataset. By confirming that these baseline algebraic elements are linearly independent over $\mathbb{Q}$ via Besicovitch's Theorem, the audit scripts analytically assess whether combination of these roots can ever mimic or cancel another. This transforms our understanding of CAPASYDIS from a multi-dimensional system that is simply observed to be "collision-free" for a specific dataset to a framework that is structurally and algebraically incapable of experiencing coordinates collisions across the entire combinatorial continuum, especially when using multiple reference points in CAPASYDIS to create multi-dimensional projections.

**2. Computational resolvability.** While Besicovitch proves that the analytical difference between two distinct sequences is strictly non-zero, we must also ensure this difference is computationally detectable. This is a classic, known challenge in computational geometry called the "Sum of Square Roots" problem, which dictates the lower bounds required to simplify nested radical expressions and evaluate sign differences (Borodin et al., 1985). Blömer (1991) provided the foundational complexity framework to calculate the minimum separation gap between such sums in polynomial time, a metric recently tightened through advanced algebraic geometry via the subspace theorem (Eisenbrand et al., 2023). These cumulative frameworks prove that the mathematical "gap" between scores remains safely within the resolution of standard 64-bit floating-point precision (which distinguishes differences down to approximately 10^{-16}, or machine epsilon). In our NR99 and M10bases analyses, we determined that a resolution filter of 10^{-10} was empirically sufficient.

#### Cited literature

Blömer, J. (1991). Computing sums of radicals in polynomial time. *Proceedings of the 32nd Annual Symposium on Foundations of Computer Science (FOCS)*, 670-677.

Borodin, A., Fagin, R., Hopcroft, J. E., & Tompa, M. (1985). Decreasing the nesting depth of expressions involving square roots. Journal of Symbolic Computation, 1(2), 169–188. https://doi.org/10.1016/s0747-7171(85)80013-4

Eisenbrand, F., Haeberle, M., & Singer, N. (2023). An improved bound on sums of square roots via the subspace theorem. arXiv. https://doi.org/10.48550/arxiv.2312.02057
